# The LINC complex ensures accurate centrosome positioning during prophase

**DOI:** 10.1101/2023.06.14.544905

**Authors:** Joana T. Lima, António J. Pereira, Jorge G. Ferreira

**Affiliations:** Instituto de Investigação e Inovação em Saúde (i3S), Porto, Portugal; Departamento de Biomedicina, Unidade de Biologia Experimental, Faculdade de Medicina do Porto, Portugal; Programa Doutoral em Biomedicina, Faculdade de Medicina, Universidade do Porto, Portugal

**Author notes:** correspondence to: Jorge G. Ferreira (ORCID: 0000-0001-6802-3696) - Address: Rua Alfredo Allen, 4200-135 Porto, Portugal.

**Keywords:** Centrosome, prophase, nuclear envelope, LINC complex, dynein

## Abstract

Accurate centrosome separation and positioning during early mitosis relies on force-generating mechanisms regulated by a combination of extracellular, cytoplasmic, and nuclear cues. The identity of the nuclear cues involved in this process remains largely unknown. Here, we investigate how the prophase nucleus contributes to centrosome positioning during the initial stages of mitosis, by using a combination of cell micropatterning, high-resolution live-cell imaging and quantitative 3D cellular reconstruction. We show that in untransformed RPE-1 cells, centrosome positioning is regulated by a nuclear signal, independently of external cues. This nuclear mechanism relies on the linker of nucleoskeleton and cytoskeleton (LINC) complex that controls the timely loading of dynein on the nuclear envelope (NE), providing spatial cues for robust centrosome positioning on the shortest nuclear axis, prior to nuclear envelope permeabilization (NEP). Our results demonstrate how nuclear-cytoskeletal coupling maintains a robust centrosome positioning mechanism to ensure efficient mitotic spindle assembly.

**Summary blurb:** Centrosome positioning in prophase is essential for efficient spindle assembly. Here, we show that centrosome positioning requires LINC complex-mediated loading of dynein on the nuclear envelope.

## Introduction

Mitosis is the process by which a cell divides its genetic content into two identical daughter cells. The start of this process is typically defined as the moment when cells start to condense their chromosomes inside the nucleus (Maddox et al. 2006; Antonin and Neumann 2016). This occurs with a near simultaneous remodelling of their cytoskeleton and a decrease in cell-matrix adhesions (Dao et al. 2009), a process regulated via integrin and cadherin signalling (Mui et al. 2016). Cytoskeletal restructuring is a consequence of the dynamic changes that occur in the microtubule (Zhai et al. 1996; Mchedlishvili et al. 2018) and actomyosin networks (Maddox and Burridge 2003; Chugh and Paluch 2018),, required to build a mitotic spindle and a stiff mitotic cortex. Subsequently, nuclear pore complexes (NPCs) (Linder et al. 2017) disassemble and the nuclear lamina depolymerizes (Heald and McKeon 1990), resulting in nuclear envelope permeabilization (NEP). These global changes are controlled by mitotic kinases such as PLK1 and CDK1 (Gavet and Pines 2010; Gheghiani et al. 2017; Jones et al. 2018), the latter ensuring coordination of cytoplasmic and nuclear events (Gavet and Pines 2010).

The goal of mitosis is to ensure the accurate segregation of chromosomes. This requires the assembly of a microtubule-based mitotic spindle that interacts with chromosomes via the kinetochores, (Toso et al. 2009; Thompson and Compton 2011). In human cells, the establishment of a bipolar spindle relies primarily on centrosomes, (Tanenbaum and Medema 2010; Agircan et al. 2014), that separate along the nuclear envelope (NE) during prophase, through forces generated by motor proteins, such as kinesin-5 and dynein (Raaijmakers et al. 2012; van Heesbeen et al. 2013). Importantly, the extent of centrosome separation and their positioning at the moment of NEP has been shown to influence mitotic fidelity (Whitehead et al. 1996; Kaseda et al. 2012; Silkworth et al. 2012; Nunes et al. 2020), and mitotic timing (Nunes et al. 2020), respectively.

Once a bipolar spindle is assembled, it must then orient inside the cell, to define a division axis. The involvement of cortical cues in mitotic spindle orientation during metaphase is well-established (Théry et al. 2005; Théry et al. 2007; Petridou and Skourides 2014). This mechanism requires the localization of the LGN-Gα1-NuMA complex to the cell cortex, which then recruits dynein to generate pulling forces on astral microtubules. However, during the early stages of mitosis this process appears to be fundamentally different. Accordingly, we have shown that during prophase, NE- associated dynein, together with Arp2/3 activity, are required to position centrosomes in the shortest nuclear axis, independently of cortical dynein (Nunes et al. 2020). These observations suggest that a nuclear cue is required for accurate centrosome positioning prior to NEP. However, the molecular nature of this nuclear cue is still unclear.

The nucleus and some of its components have already been proposed to impact centrosome function. Accordingly, perturbations in the nuclear lamina due to loss of lamin A, leads to impaired centrosome separation and asymmetric NPC distribution in prophase (Guo and Zheng 2015; Boudreau et al. 2019). In addition, different components of the linker of nucleoskeleton and cytoskeleton (LINC) complex, required for nucleo-cytoplasmic coupling (Lombardi et al. 2011), have been implicated in centrosome-nucleus tethering during cell migration, by directly associating with NE- bound dynein (Malone et al. 2003; Zhang et al. 2009; Splinter et al. 2010; Boudreau et al. 2019). Moreover, the LINC complex also facilitates centrosome separation during prophase (Stiff et al. 2020) and decreases chromosome scattering during prometaphase, aiding their capture after NEP (Booth et al. 2019). Nevertheless, how these cytoplasmic and nuclear players interact during the G2/M transition to ensure accurate centrosome positioning and efficient spindle assembly remains unknown. Here, we identify the LINC complex as essential for defining the centrosome-nuclear axis during prophase in normal, near-diploid cells, by orienting centrosomes towards the shortest nuclear axis.

## Results

### Chromosomally stable RPE-1 cells systematically position their centrosomes at the shortest nuclear axis

It has been previously shown that centrosome separation and positioning at the moment of NEP impacts mitotic progression and fidelity (Kaseda et al. 2012; Silkworth et al. 2012; Nunes et al. 2020). However, the degree of conservation of this mechanism between normal, untransformed cells and cancer cell lines is still unknown. To address this, we performed live-cell imaging of chromosomally stable RPE-1 cell line and two chromosomally unstable cancer cell lines of different origins (U2-OS derived from osteosarcoma and MDA-MB-468 derived from a breast adenocarcinoma), with high spatial and temporal resolution. All cell lines were seeded on micropatterned lines with a 10μm width, coated with fibronectin (FBN; Fig. 1A). This allowed us to accurately standardise cell shape and intracellular organization(Théry et al. 2005; Nunes et al. 2020). Then, we reconstructed the dynamics of centrosomes, nucleus, and cell membrane during the G2-M transition, using our previously developed tool, Trackosome (Castro et al. 2020) (Fig. S1A-E). With this approach, we can obtain quantitative data and correlate several parameters. Namely, we determined the alignment of the longest cell axis with the longest nuclear axis (Fig. S1F), correlated the alignment of a vector connecting the two centrosomes with the longest nuclear axis (angle centrosome – nucleus, Fig. S1G) and measured the separation of the two centrosomes to opposite sides of the nucleus, by determining the angle formed by a vector connecting the two centrosomes that intercepts the nuclear centroid (angle centrosome – centrosome, Fig. S1H). This allowed us to construct polar plots that display the alignment of the centrosomes with the longest nuclear axis (Fig. 1SI; note that an orientation towards 90° corresponds to alignment with the shortest nuclear axis) and graphs that correlate centrosome positioning and separation to opposite sides of the nucleus (Fig. S1J).

**Figure 1.**
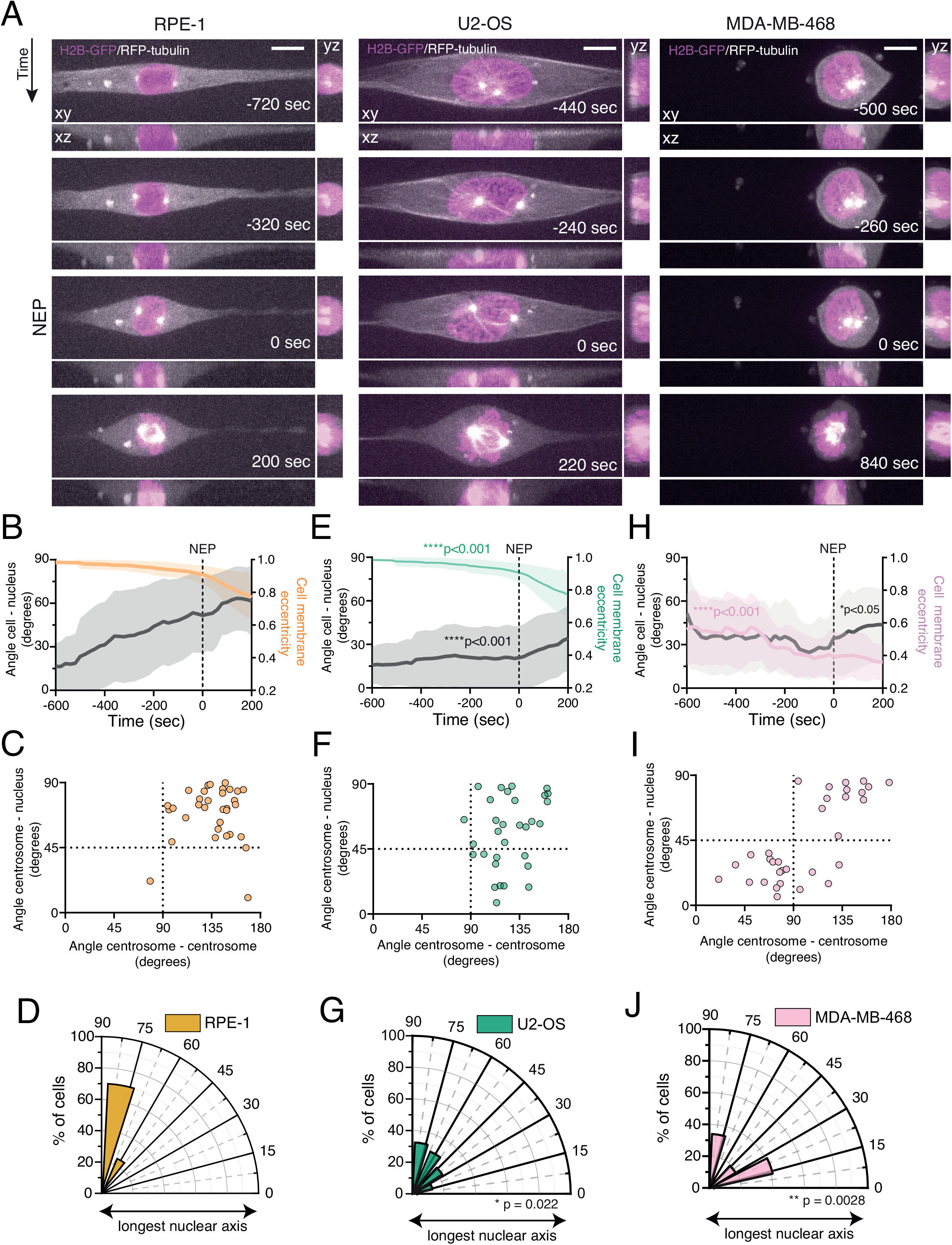
Chromosomally stable, near-diploid cells efficiently separate and position their centrosomes at the shortest nuclear axis. **(A)** Representative frames from a movie of a near-diploid, untransformed RPE-1 cell (left panel) and chromosomally unstable U2-OS (middle panel) and MDA-MB-486 (right panel) cancer cells, stably expressing H2B-GFP and RFP-tubulin, seeded on a 10μm wide-micropatterned line, showing centrosome movement and the overall cellular reorganisation in preparation for mitotic entry. Time is in sec and time zero corresponds to NEP. Scale bar = 10μm. **(B)** Quantification of cell membrane eccentricity (orange) and the angle between the longest cell axis and longest nuclear axis (angle cell- nucleus; gray), of RPE-1 cells during mitotic entry, obtained with our custom-made MATLAB script. Line represents mean value and shaded area corresponds to SD (n=33). Dashed line represents the moment of NEP. **(C)** Correlation between centrosome separation to opposite sides of the nucleus (angle centrosome-centrosome; x axis) and centrosome positioning relative to the longest nuclear axis (angle centrosome-nucleus; y axis), at the moment of NEP for RPE-1 cells. Cells that efficiently separate their centrosomes will present high values of centrosome-centrosome angle, and therefore locate at the right half of the graph. Cells that correctly position their centrosomes at the shortest nuclear axis will cluster at the upper half of the graph. **(D)** Polar plot quantifying centrosome positioning relative to the longest nuclear axis at NEP for RPE-1 cells. **(E)** Quantification of cell membrane eccentricity (green; ****p<0.001) and angle cell-nucleus (gray; ****p<0.0001) for U2-OS cells (n=31). **(F)** Correlation between the angle centrosome-centrosome (*p=0.036) and the angle centrosome- nucleus for U2-OS cells. **(G)** Polar plot quantifying centrosome positioning relative to the longest nuclear axis at NEP for U2-OS cells (*p= 0.022). **(H)** Quantifications of cell membrane eccentricity (pink; ****p<0.0001) and angle cell-nucleus (gray; * p=0.011) for MDA-MB-468 cells (n=32). **(I)** Correlation between the angle centrosome-centrosome (****p<0.001) and the angle centrosome-nucleus, for MDA-MB-468 cells. **(J)** Polar plot quantifying centrosome positioning relative to the longest nuclear axis at NEP for these MDA-MB-468 cells (**p=0.028).

Detailed analysis of RPE-1 cells during the G2-M transition revealed that the long axis of the nucleus is initially aligned with the long cell axis (Fig. 1A, left panel and 1B; movie S1), likely due to a geometrical constraint imposed by the line micropattern, as was previously shown for interphase cells (Versaevel et al. 2012). Once cells start to round up in preparation for mitosis, as measured by the decrease in cell membrane eccentricity, the long nuclear axis progressively diverges from the long cell axis (Fig. 1B), suggesting that mitotic cell rounding uncouples the two parameters. This reorientation occurs simultaneously with the movement of centrosomes to opposite sides of the nucleus (Fig 1C). Consequently, at NEP, centrosomes are positioned on the shortest nuclear axis (Fig. 1C, 1D). Importantly, this behaviour is not observed in the two cancer cell lines analysed. In U2-OS cells, mitotic cell rounding is altered in comparison with RPE-1 cells (Fig. 1A, middle panel; 1B, 1E; ****p<0.001) and the nucleus maintains alignment with the main cell axis almost until the moment of NEP (Fig. 1E; movie S2; ****p<0.001). As a result of this physical constraint, centrosomes separate, but fail to position on the shortest nuclear axis at NEP (Fig. 1F, 1G; *p=0.022). On the other hand, MDA-MB-468 cells are morphologically distinct from RPE-1 or U2- OS. They are rounder and have a smaller area of adhesion to the micropatterned substrate (Fig. 1A, right panel; movie S3). As a result, they do not exhibit the decrease in cell membrane eccentricity normally observed during mitotic cell rounding (Fig. 1B, 1H; ****p<0.001). Consequently, the long nuclear axis is randomly oriented inside the cell (Fig. 1H; p<0.05). In addition, centrosomes in MDA-MB-468 often show incomplete separation and are unable to reach opposite sides of the nucleus at NEP (Fig. 1I). This defect in centrosome separation could be due to the decrease in cell adhesion, which affects the activity of kinesin-5 (Nunes et al. 2020; Kamranvar et al. 2022), essential for the initial stages of centrosome separation. Consequently, centrosomes are randomly positioned relative to the nucleus at the moment of NEP (Fig. 1J; **p=0.0028). Overall, our data show that RPE-1 cells have a robust centrosome positioning mechanism during the G2-M transition, and this mechanism is compromised both in U2-OS and MDA-MB-468 cells.

### Mitotic cell rounding is not the major determinant for centrosome positioning

Given that the U2-OS and MDA-MB-468 cell lines that we tested failed to correctly position centrosomes on the shortest nuclear axis and displayed altered mitotic cell rounding, we wondered if the two events were interconnected. This is particularly relevant since it was previously shown that mitotic rounding is essential for providing the necessary space to assemble a mitotic spindle (Lancaster et al. 2013; Sorce et al. 2015; Dix et al. 2018; Nunes et al. 2020). To test this hypothesis, we proceeded to interfere with mitotic cell rounding in RPE-1 cells, by either impairing it or accelerating it. Mitotic rounding requires both cortical retraction and adhesion disassembly. Therefore, we started by acutely treating cells with a Rho-associated protein kinase (ROCK) inhibitor (Y-27632; Fig. 2A, B), known to decrease actomyosin contractility and delay cortical retraction (Maddox and Burridge 2003). Upon treatment with Y27632, centrosomes were still able to separate and position correctly on the shortest nuclear axis (Fig. 2C-F), although the rate of cell rounding was delayed due to an impairment of cortical retraction (Fig. 2G, ***p<0.001). This is in agreement with our previous observations in HeLa cells (Nunes et al. 2020). Next, we proceeded to interfere with adhesion disassembly by expressing a mutant form of Rap1 (Rap1Q63E; Rap1*), that effectively blocks mitotic rounding (Dao et al. 2009) (Fig. 2H-J, ***p<0.001). Delaying adhesion disassembly in RPE-1 cells did not affect centrosome separation or positioning on the shortest nuclear axis (Figure 2K-N; p=0.583), indicating that cell rounding impairment in RPE-1 cells does not significantly impact centrosome positioning at the moment of NEP.

**Figure 2.**
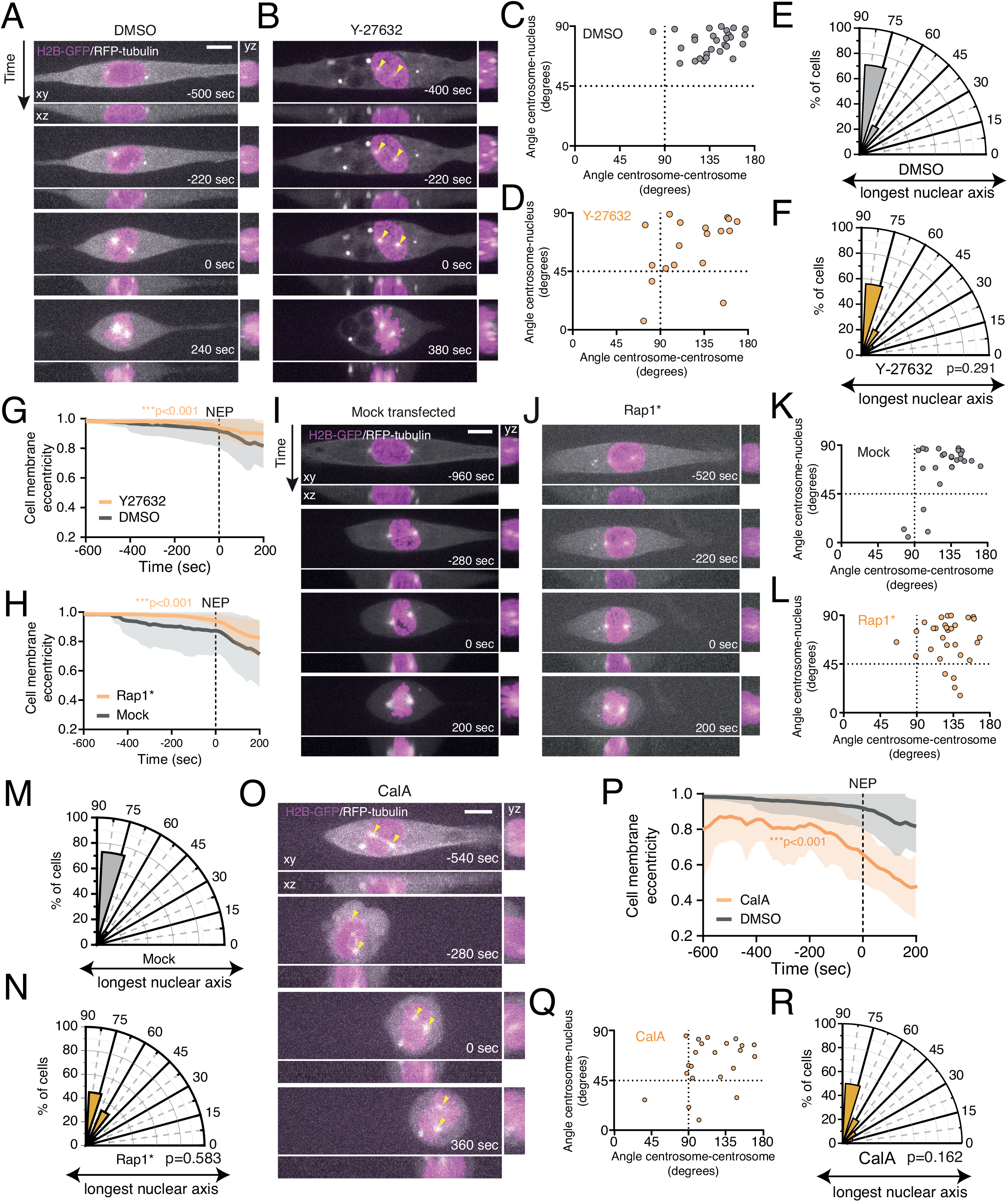
Cell rounding status does not influence centrosome positioning in RPE-1 cells. Representative frames from a movie of an RPE-1 cell stably expressing H2B-GFP and RFP-tubulin, seeded on a 10μm wide-micropatterned line, treated with **(A)** DMSO and Y-27632, showing centrosome movement. Yellow arrowheads indicate centrosome position. Correlation between centrosome separation (angle centrosome-centrosome; x axis) and centrosome positioning (angle centrosome-nucleus; y axis;), at the moment of NEP, for DMSO-treated (**C**; n=28) or Y-27632-treated RPE-1 cells (**D**; n=18). Polar plot quantifying centrosome positioning relative to the longest nuclear axis at NEP for these RPE-1 cells treated with DMSO (**E**) or Y-27632 (**F**; p=0.291). Quantification of cell membrane eccentricity of DMSO-treated cells (**G**; gray) and Y-27632-treated cells (orange; ****p<0.001) or mock transfected (**H**; Mock; gray; n=25) and Rap1* transfected cells (orange; n=31; ****p<0.001). Representative frames from movies of mock (**I**) and Rap1* (**J**) transfected RPE-1 cells, stably expressing H2B-GFP and RFP-tubulin, seeded on a 10μm wide-micropatterned line. Correlation between the angle centrosome-centrosome (x axis) and the angle centrosome-nucleus (y axis), at the moment of NEP for Mock (**K**; n=26) and Rap1* (**L**; n=31) transfected cells. Polar plot quantifying centrosome positioning relative to the longest nuclear axis at NEP for Mock (**M**) or Rap1* (**N**; p=0.583) transfected RPE-1 cells. (**O**) Frames from a representative movie of a CalyculinA (CalA) treated RPE-1 cell, stably expressing H2B-GFP and RFP- tubulin, seeded on a 10μm wide-micropatterned line. (**P**) Cell membrane eccentricity of CalA-treated cells (orange; n=22; ****p<0.001). (**Q**) Correlation between the angle centrosome-centrosome (x axis) and the angle centrosome-nucleus (y axis) for cells treated with CalA. (**R**) Polar plot quantifying centrosome positioning relative to the longest nuclear axis at NEP for cells treated with CalA (p=0.162). For all movies, time is in sec. and time zero corresponds to NEP. Scale bars, 10μm.

Next, we tested the effect of inducing premature rounding by treating cells with Calyculin-A (CalA), a protein phosphatase inhibitor that increases the phosphorylation of Ezrin/Radixin/Moesin (ERM) proteins and triggers cell rounding in adherent cells (Tachibana et al. 2015). Acute treatment with CalA led to a faster rounding of the cells (Fig. 2O, P; ***p<0.001), as anticipated. Nevertheless, this did not significantly affect centrosome separation or positioning at NEP (Fig. 2Q, R; p=0.162). Overall, we conclude that centrosome positioning in RPE-1 cells during early mitosis is a robust process, that is largely independent of mitotic cell rounding.

To further investigate the relevance of cell rounding for centrosome positioning, we then decided to interfere with mitotic cell rounding in the U2-OS and MDA-MB-468 cancer cell lines. We either forced adhesion in the highly rounded MDA-MB-468 or promoted rounding in the flatter U2-OS cells. We started by transfecting MDA-MB-468 cells with

Rap1*. As expected, Rap1* expression delayed cell rounding (Fig. S2A, B; ****p<0.001), but this did not restore correct centrosome separation or positioning (Fig. S2C, D). Next, we tried to interfere with rounding in U2-OS cells by using Y-27632 or CalA (Fig. S2E-I). Neither treatment was sufficient to restore centrosome positioning on the shortest nuclear axis (Fig. S2J-M). Because these results were obtained from cells seeded also wanted to rule out the possibility that the failure to position centrosomes could be due to geometrical constraints imposed by the micropattern. Therefore, we seeded U2-OS cells on micropatterned lines with 20μm width or on non-patterned FBN-coated coverslips. In all these conditions, U2-OS cells were still unable to correctly position their centrosomes at the shortest nuclear axis (Fig. S2N-S), even though they showed a significantly decrease in cell membrane eccentricity when placed in non-patterned substrates, reflecting a more rounded state (Fig. S2T; ****p<0.0001). Taken together, our results indicate that centrosome positioning on the shortest nuclear axis in normal, untransformed RPE-1 cells is independent of the rounding state of the cell and that the defects in positioning observed in the U2-OS and MDA-MB-468 cancer cells cannot be rescued by manipulating cell rounding.

### The nuclear lamina is not required for centrosome positioning on the shortest nuclear axis

Given these results and our previous observations (Nunes et al. 2020), it seems unlikely that an external cue could be driving centrosome positioning during early mitosis. Therefore, we hypothesized that a nuclear cue could be responsible for ensuring accurate centrosome positioning on the shortest nuclear axis. One possible candidate is the nuclear lamina, since it was previously shown that lamins can position NPCs and centrosomes (Guo and Zheng 2015) and are required for centrosome separation (Boudreau et al. 2019). We started by analysing the levels of nuclear lamina components in each of the cell lines in our panel, by measuring the fluorescence intensity of lamin A/C and lamin B1 at the NE (Fig. S3A, B). We observed that the levels of both Lamin A/C and Lamin B1 were altered in U2-OS and MDA-MB-468, when compared to RPE-1 cells (Fig. S3C, D; ****p<0.001). Consequently, the ratios of lamin A/C relative to lamin B1 in the two cancer cell lines were substantially lower than in RPE-1 cells (Fig. S3E). Our results are in agreement with lamin expression data available from the online repository Cancer Dependency Map Project (DepMap) (Fig. S3F). Moreover, the decrease in lamin A levels observed in cancer cells was accompanied by a decrease in nuclear solidity (Fig. S3G, **** p<0.0001), which suggests that nuclear structure in both U2-OS and MDA-MB-468 cells is altered when compared to RPE-1 cells. Thus, we decided to manipulate lamin levels in RPE-1 cells and ask whether this was sufficient to impair centrosome positioning on the shortest nuclear axis. We started by depleting Lamin A in RPE-1 cells, using RNA interference (siLMNA; Fig. 3A-C). However, this depletion was not sufficient to alter centrosome behaviour, as cells still efficiently separated and positioned them at the shortest nuclear axis (Fig. 3D-G). To confirm our observations, we next sought to imbalance the lamin A:B ratio by overexpressing a HaloTag9-Lamin B1 construct in RPE-1 cells (Fig. 3H, I and Fig. S3H; ****p<0.0001). Similarly to Lamin A depletion, Lamin B1 overexpression also did not disrupt correct centrosome positioning at NEP (Figure 3J-M). Taken together, these results indicate that even though the ratio of lamin A:B is perturbed in the two cancer cells used for this study, this is likely not the cause for the observed errors in centrosome positioning at NEP, as experimental manipulation of the levels of either lamin A or B in RPE-1 cells did not affect this process. Next, we sought to interfere with additional nuclear components, independently of lamin levels. To do so, we overexpressed an mCherry-tagged version of lamin B receptor (LBR), an integral protein of the inner nuclear membrane (INM) that associates with the nuclear lamina (Fig. S3I; ****p<0.0001) (Appelbaum et al. 1990). It has been previously shown that LBR overexpression causes perinuclear ER expansion and the overproduction of nuclear membranes (Ma et al. 2007), leading to NE folding and altered nuclear structure (Dantas et al. 2022), similar to what we observed in U2-OS and MDA-MB-468 cells. However, interfering with nuclear structure by overexpressing LBR in RPE-1 cells (Fig. 3 N, O), did not affect the separation and positioning of centrosomes on the shortest nuclear axis (Fig. 3P-S). Overall, our data indicates that neither the nuclear lamina nor LBR are required for correct centrosome positioning during early mitosis. They further suggest that the altered lamin levels in U2-OS and MDA-MB-468 are likely not the cause for the mispositioned centrosomes observed in these cell lines.

**Figure 3.**
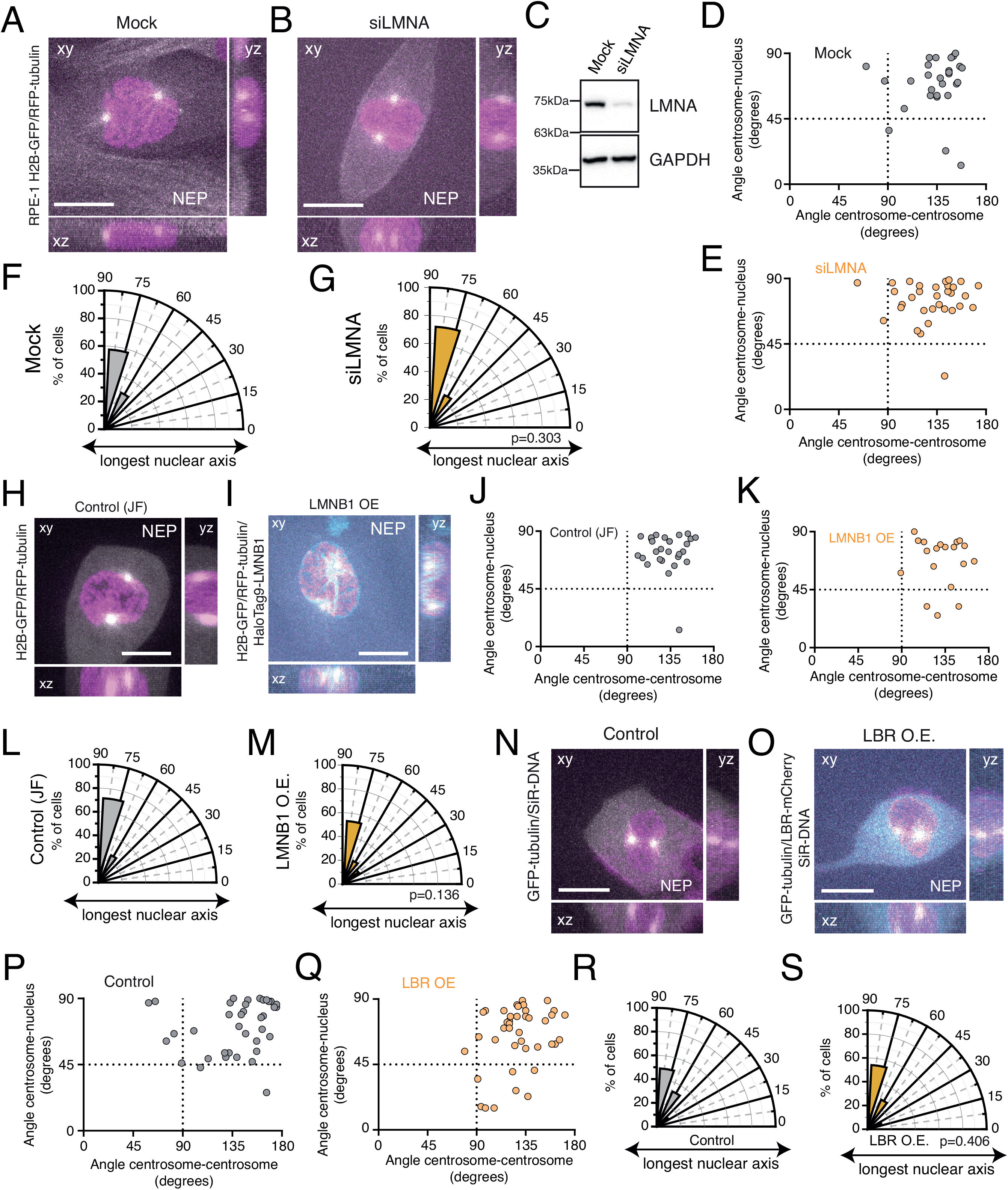
Nuclear lamina components do not affect centrosome positioning. Representative frames from movies of Mock (**A**) or Lamin A depleted (**B**; siLMNA) RPE-1 cells at the moment of NEP, stably expressing H2B-GFP and RFP-tubulin. (**C**) Western blot showing depletion efficiency for these cells. Correlation between the angle centrosome-centrosome (x axis) and the angle centrosome-nucleus (y axis) at the moment of NEP, for Mock (**D**; n=28) and Lamin A depleted (**E**; n= 32) cells. Polar plots quantifying centrosome positioning relative to the longest nuclear axis at NEP for Mock (**F**) and LMNA depleted RPE-1 cells (**G**; p=0.303). Representative frame of the moment of NEP from movies of RPE-1 cells stably expressing H2B-GFP and RFP-tubulin (**H**; control treated with JF647; n=28) or overexpressing HaloTag9-LMNB1 (**I**; LMNB1 OE; n=19). Correlation between the angle centrosome-centrosome (x axis) and the angle centrosome-nucleus (y axis; p=0.216) at the moment of NEP, for Control cells (**J**) and cells overexpressing LMNB1 (**K**). Polar plots quantifying centrosome positioning relative to the longest nuclear axis at NEP for Control (**L**) and LMNB1 overexpressing RPE-1 cells (**M**; p=0.136). Representative frames of the moment of NEP from movies of Control (**N**; n=37) or LBR-mCherry overexpressing (**O**; n=41) RPE-1 cells, stably expressing GFP-tubulin and treated with SiR-DNA, plated on fibronectin. Correlation between the angle centrosome-centrosome and the angle centrosome-nucleus at the moment of NEP, for Control (**P**) and LBR overexpressing (**Q**) cells. Polar plots quantifying centrosome positioning relative to the longest nuclear axis at NEP for Control (**R**) and LBR overexpressing RPE-1 cells (**S**; p=0.406). For all images, scale bars = 10μm.

### The LINC complex is required for centrosome positioning on the shortest nuclear axis

Next, we focused our attention on the LINC complex, since it was previously shown to play a role in early mitosis (Booth et al. 2019; Stiff et al. 2020). Immunolocalization of SUN1 and SUN2 in prophase cells revealed that both proteins localized to the NE, as expected (Fig. 4A, B). Interestingly, in a significant proportion of cells, SUN1 and SUN2 also concentrated in the area between and around the two separating centrosomes and their corresponding microtubule arrays (Fig. 4A, B, insets). This localization was dependent on microtubules, since treatment with low doses of nocodazole (Noc) disrupted inter-centrosomal SUN localization (Fig. 4C, D). We then proceeded by depleting individual SUN proteins (SUN1 and SUN2), in RPE-1 cells using a lentiviral-mediated short hairpin RNA (shRNA), giving rise to a heterogeneous population of cells with different levels of depletion (Fig. 4A, B and Fig. S4A-D). This was sufficient to decrease NE localization and inter-centrosomal accumulation of SUN1 and SUN2 (Fig. 4A, B). We then assessed the impact of SUN1 and SUN2 depletion for centrosome behaviour throughout mitotic entry. Upon depletion, these cells were still capable of efficiently separating centrosomes (Figure 4E-J; Sup. movies S4 and S5). However, they had an impaired positioning of centrosomes at the shortest nuclear axis (Figure 4K-M; *p=0.0165 and p=0.0175 for SUN1 and SUN2, respectively).

**Figure 4.**
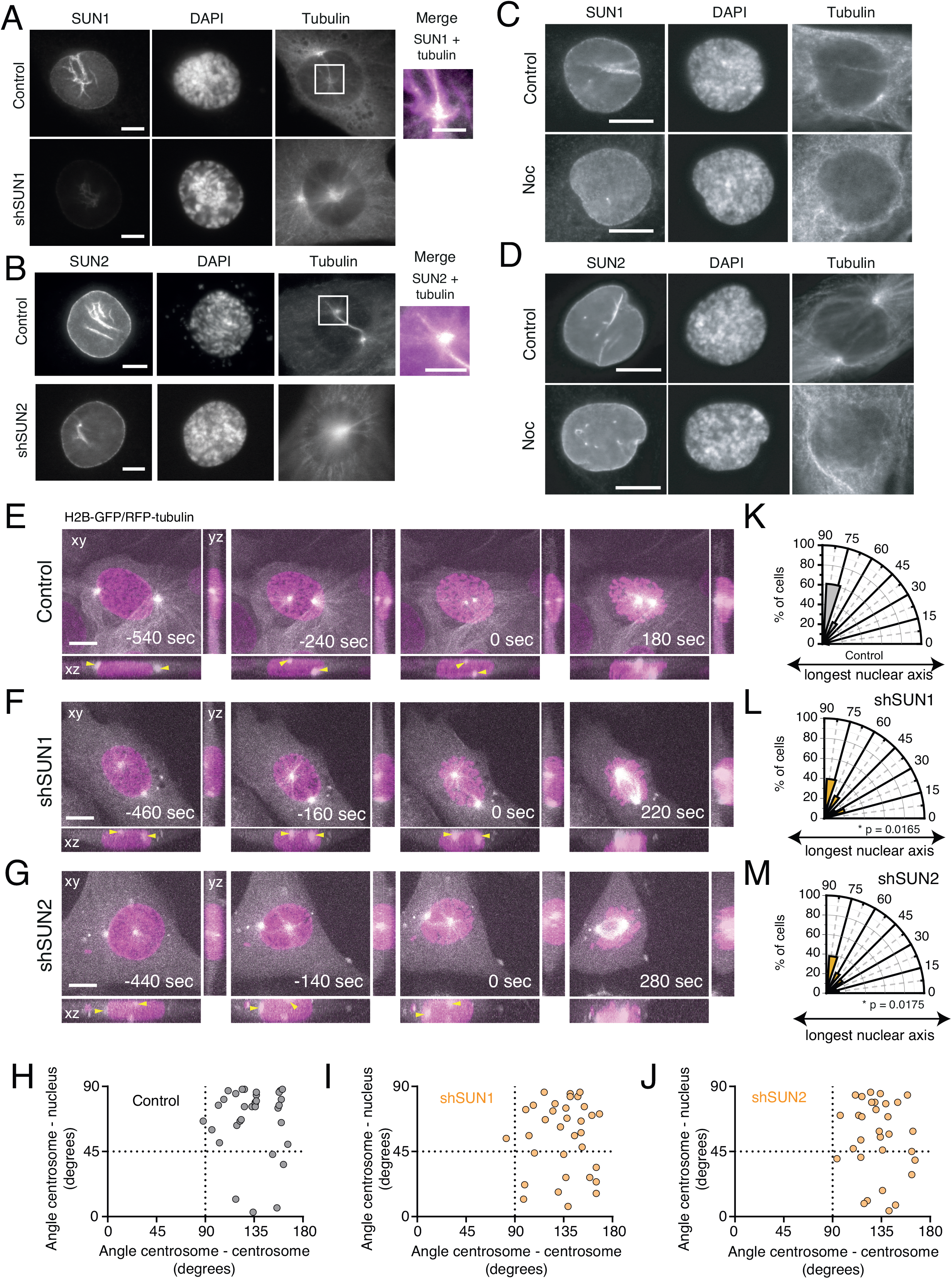
SUN proteins are required for correct centrosome positioning. (**A**) Representative immunofluorescence images of Control and shSUN1 treated cells, immunostained for SUN1. (**B**) Representative immunofluorescence images of Control and shSUN2 treated cells cells, immunostained for SUN2. Note the co-localisation between tubulin and the SUN proteins in the merged image inset. Representative immunofluorescence images of Control and shSUN1-treated RPE-1 cells (**C**) or Control and shSUN2-treated cells (**D**). Nocodozole (Noc) was added to the cells for 10 minutes, to induce microtubule depolymerisation. Note the decrease in SUN1 and SUN2 staining between the centrosomes. Representative frames from movies of Control (**E**), shSUN1- treated (**F**) or shSUN2-treated (**G**) cells, stably expressing H2B-GFP and RFP-tubulin, seeded on fibronectin, during mitotic entry. Time is in sec. Time zero corresponds to NEP. Yellow arrows indicate centrosomes position. Correlation between the angle centrosome-centrosome (x axis) and the angle centrosome-nucleus (y axis) at the moment of NEP, for Control cells (**H**; n= 33), shSUN1-treated (**I**; n= 33), and shSUN2- treated (**J**; n= 31) cells. Polar plots quantifying centrosome positioning relative to the longest nuclear axis at NEP for Control (**K**), shSUN1-treated (**L**; *p=0.0165), and shSUN2-treated (**M**; * p=0.0175) cells. Yellow arrowheads indicate centrosome positions. All scale bars, 10μm.

Given the results obtained with SUN1 and SUN2, we then tested whether nesprins, which are part of the LINC complex and interact with SUN proteins in the perinuclear space, could also impact centrosome function. For that purpose, we expressed a dominant-negative form of KASH tagged with mCherry (DN-KASH; Fig. S4E; Sup. Movie S6 and S7) that prevents binding of endogenous nesprins to SUN proteins and blocks force propagation across the NE (Lombardi and Lammerding 2011; Zhang et al. 2019). As a corresponding control, we generated a cell line expressing a mCherry-tagged mutant version of the same DN-KASH construct where the last 4 aminoacids (PPPL) of the KASH domain were removed (ΔPPPL-KASH). This prevents ΔPPPL-KASH from interacting with SUN1 or SUN2, thus making it a “defective” dominant negative (Zhang et al. 2019). Firstly, we confirmed by immunofluorescence the ability of the DN-KASH construct displace endogenous nesprins from the NE, as assessed by nesprin 2 localization (Fig. 5A; SYNE-2). Contrarily, and as expected, expression of the ΔPPPL-KASH construct did not interfere with nesprins localization on the NE, similarly to unmanipulated RPE-1 cells (Fig. 5A and S4F). We then analysed centrosome behaviour in these cells. Upon expression of ΔPPPL-KASH, cells behaved similarly to unmanipulated RPE-1 cells, with an efficient separation and positioning of centrosomes on the shortest nuclear axis (Fig. 1C, 5D, F; p=0.708 for ΔPPPL-KASH compared with unmanipulated RPE-1 cells). However, DN-KASH-expressing cells showed compromised separation and positioning of centrosome (Fig. 5E, G; *p=0.0155 and *p=0.0237, respectively).

**Figure 5.**
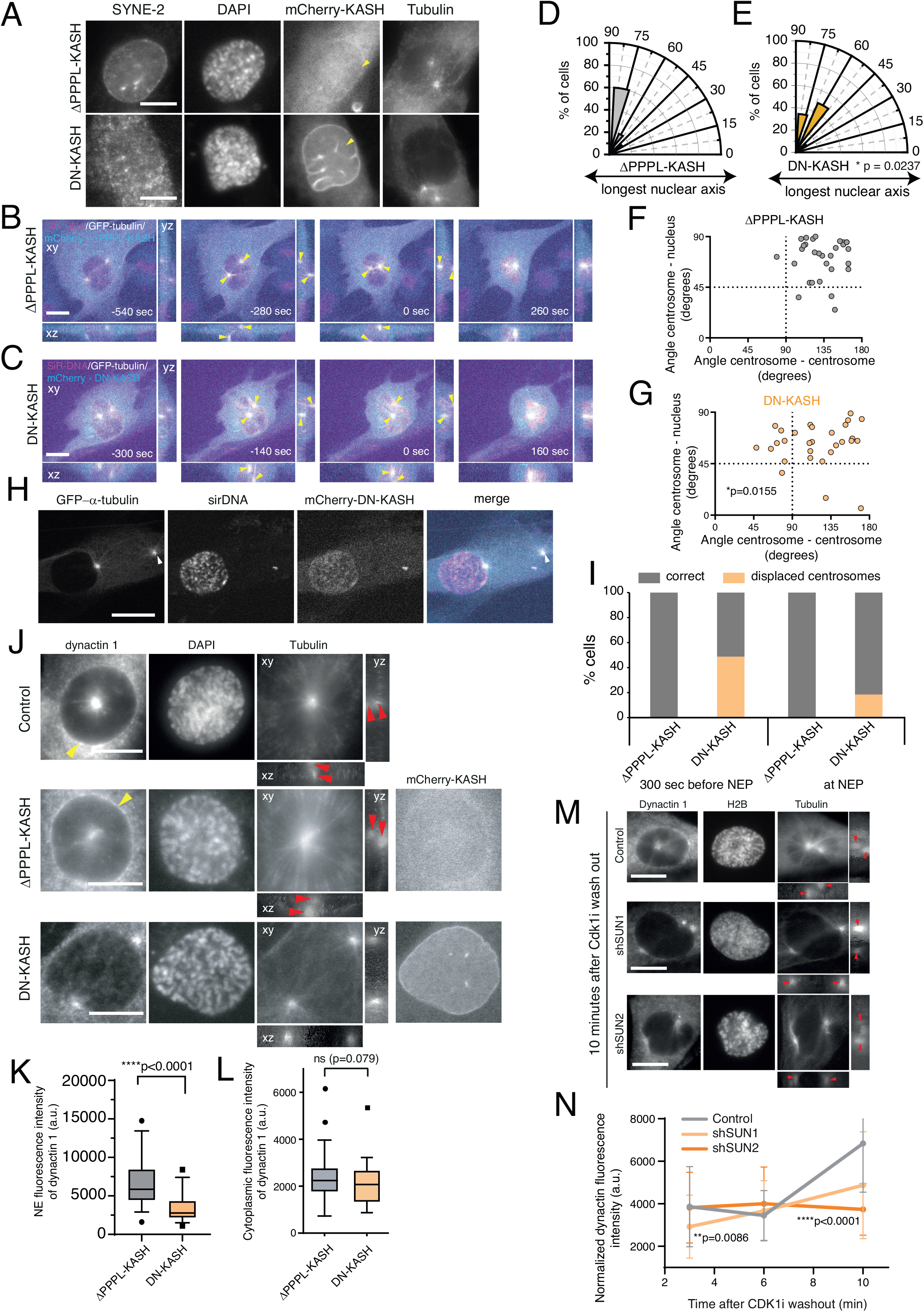
The LINC complex is required for centrosome positioning on the shortest nuclear axis. **(A)** Representative immunofluorescence images of a ΔPPPL-KASH (top panel) and a DN-KASH (bottom panel) cell, immunostained for nesprin-2 (SYNE-2). Note how expression of DN-KASH displaces nesprin-2 from the NE (yellow arrows), as opposed to expression of ΔPPPL-KASH. Representative frames from movies of Control (**B**; ΔPPPL-KASH) and DN-KASH (**C**) treated cells, stably expressing tubulin-GFP and treated with SiR-DNA, seeded on fibronectin, during mitotic entry. Yellow arrowheads indicate centrosomes position. Polar plots quantifying centrosome positioning relative to the longest nuclear axis at NEP for RPE-1 expressing ΔPPPL-KASH (**D**; n= 30) or DN- KASH cells (**E**; n= 29; *p=0.0237). Correlation between the angle centrosome- centrosome (x axis) and the angle centrosome-nucleus (y axis) at the moment of NEP for Control (**F**; ΔPPPL-KASH) or DN-KASH expressing (**G**) cells. (**H**) Representative frame from a movie of a DN-KASH expressing cell, stably expressing Tubulin-GFP and treated with SiR-DNA, highlighting centrosome detachment from the NE (white arrowheads). (**I**) Quantification of centrosome displacement from the NE during mitotic entry for ΔPPPL-KASH or DN-KASH expressing cells before (-300 sec) and at NEP. (**J**) Representative immunofluorescence images of control (top panel), ΔPPPL-KASH (middle panel) or DN-KASH (bottom panel) cells, labelled for the dynein adaptor, dynactin1. Please note the decreased dynactin1 signal at the NE of DN-KASH expressing cells, when compared to controls or ΔPPPL-KASH cells (yellow arrowheads). Red arrowheads indicate centrosomes positioned above and beneath the nucleus (lateral projections). Quantification of the normalized fluorescence intensity signal of dynactin 1 signal on the NE (**K**; ***p<0.001) and cytoplasm (**L**; p=0.079) of ΔPPPL-KASH (n= 36) or DN-KASH (n= 35) cells. Time is in sec and time zero corresponds to NEP. Scale bars, 10μm.

Finally, we decided to test whether the cancer cell lines used in this study have an altered LINC complex, considering that they show centrosome positioning defects (Fig. S5). To, we achieve this, quantified the levels of different LINC complex components on the NE by performing immunofluorescence to detect SUN1, SUN2, SYNE1 (nesprin 1) and SYNE2 (nesprin 2). Our results indicate that U2-OS cells have a significant decrease in the levels of nesprin 1 and SUN2 (Fig. S5A, B, E; ****p<0.0001), whereas MDA-MB-468 show a decrease of nesprin1, SUN1 and SUN2 (Fig. S5A, B, D, E: ****p<0.001; **p<0.01). Intriguingly, both cancer cell lines show increased levels of nesprin 2, when compared to RPE-1 (Fig. S5C; ****p<0.001). Nevertheless, these data clearly indicate that the LINC complex is altered in cancer cells, which correlates with their inability to correctly position centrosomes. Overall, our data strongly suggest that the LINC complex provides the cues for positioning centrosomes on the shortest nuclear axis during prophase.

### Dynein recruitment to the NE during early mitosis requires a functional LINC complex

Next, we sought to determine how LINC complex disruption affected centrosome positioning during prophase. While imaging DN-KASH-expressing cells, we frequently observed centrosome detachment from the NE during earlier stages of prophase (Fig. 5H, 5I), which was never observed in cells expressing the ΔPPPL-KASH construct (Fig. 5I) or unmanipulated RPE-1 cells. Curiously, this centrosome displacement from the NE decreased as cells entered mitosis (Fig. 5I). These observations suggest that disruption of the LINC complex leads to a transient defect in centrosome-NE tethering during the G2-M transition, which is rescued as cells approach NEP. Previous studies have shown that a specific pool of NE-bound dynein, is sufficient to tether centrosomes to the NE during prophase, by generating pulling forces on microtubules (Gönczy et al. 1999; Robinson et al. 1999), in a BicD2 (Splinter et al. 2010) or Nup133-dependent manner (Bolhy et al. 2011). Moreover, the LINC complex was previously shown to help maintain the centrosome-nucleus connection during neuronal migration in mice (Zhang et al. 2009). Thus, we decided to analyse whether the centrosome displacement we observed in DN-KASH cells could be due to a defective loading of dynein on the NE, triggered by a disruption of the LINC complex. For this purpose, we assessed the localization of dynactin 1, a dynein adaptor that also localizes to the NE (Splinter et al. 2012), upon expression of ΔPPPL-KASH or DN-KASH. Importantly, expression of DN-KASH significantly decreased the levels of dynactin 1 on the NE, when compared to cells expressing ΔPPPL-KASH (Fig. 5J, K; ****p<0.0001), while having no effect on the overall levels of cytoplasmic dynactin 1 (Fig. 5L; n.s. p=0.079) nor significantly changing the levels of another NE protein such as Lamin B (Fig. S4G, H; p=0.0823). To further confirm the LINC complex requirement for dynactin loading on the NE, we proceeded to analyse its levels following SUN depletion (Fig. 5M). We synchronized RPE-1 cells in late G2 using a CDK1 inhibitor (CDK1i; Ro-3306). Then, following inhibitor washout, we quantified the levels of dynactin on the NE at 3, 6 and 10 minutes, using immunofluorescence analysis (Fig. 5N). As anticipated, cells depleted of SUN1 had decreased levels of dynactin 3 min after release (**p=0.0086). On the other hand, SUN2 depleted cells had significantly lower levels of dynactin at 10 min after release (****p<0.0001). It should be noted that the effect of depleting individual SUN proteins on NE dynactin levels is not as severe as DN-KASH expression. This likely happens because expression of the DN- KASH completely disrupts the interaction with both SUN proteins at the same time. On the other hand, with individual SUN depletion, the remaining SUN protein might still associate with nesprins to maintain a partial function. Interestingly, decreased levels of NE-associated dynactin 1 were also seen in both U2- OS and MDA-MB-468 cell lines when compared to RPE-1 cells (Fig. S5F, G; ****p<0.0001), which fits nicely with the decreased levels of LINC complex components (Fig. S5A-E) and the defect in centrosome positioning (Fig. 1) exhibited by these cancer cell lines. Taken together, our results indicate that an intact LINC complex is required for dynein localization at the NE during the G2-M transition, which then allows accurate centrosome positioning at NEP.

## DISCUSSION

The manner in which centrosomes separate and position during prophase has direct implications for the efficiency of mitosis (Kaseda et al. 2012; Silkworth et al. 2012; Stiff et al. 2020). We previously reported that, during the G2-M transition, separating centrosomes exhibit a coordinated motion along the NE, so that they are positioned on the shortest nuclear axis at the moment of NEP (Nunes et al. 2020). This centrosome-nucleus configuration occurs independently of the cortical cues that dictate spindle orientation in later stages of mitosis, since these cortical complexes are only assembled after NEP (Kiyomitsu and Cheeseman 2012; Kotak et al. 2012). Importantly, this prophase configuration suggested that a nuclear signal could be providing the cues to regulate centrosome positioning during the early stages of mitosis. However, the molecular mechanism remained unclear. Here, we show that this centrosome-nuclear axis orientation is a robust, LINC complex-dependent process in near-diploid, untransformed RPE-1 cells and that this mechanism is disrupted in U2-OS and MDA-MB-468 cell lines (Fig. 6).

**Figure 6.**
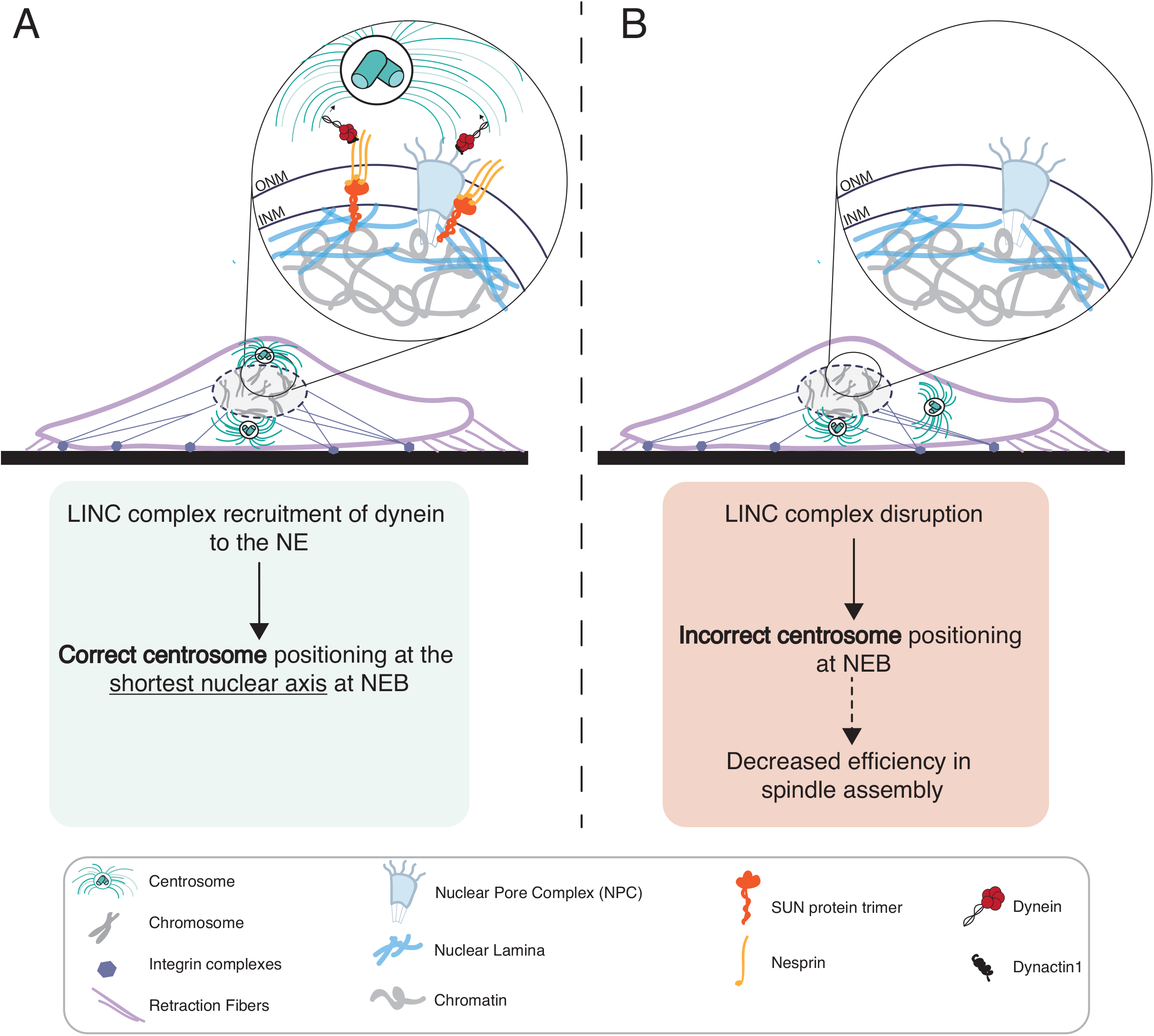
Proposed model for LINC complex-dependent centrosome positioning. **(A)** In normal conditions, the LINC complex allows timely loading of dynein on the NE, leading to correct centrosome positioning. This allows efficient mitotic spindle assembly and chromosome segregation. (B) Upon LINC complex disruption, loading of dynein on the NE is delayed, leading to incorrect centrosome positioning. Consequently, mitotic spindle assembly is perturbed, which could result in chromosomal instability (CIN), under the appropriate genetic background.

In preparation for mitosis, most cells round up and adopt a spherical shape (Taubenberger et al. 2020). This event, driven by a combination of adhesion disassembly (Dao et al. 2009) and membrane retraction (Maddox and Burridge 2003), is required to release the geometric constraints imposed by cell shape (Versaevel et al. 2012; Lancaster et al. 2013). In addition, this allows nuclear rotation during prophase (Nunes et al. 2020) and provides sufficient space for mitotic spindle assembly to occur without errors (Tse et al. 2012; Lancaster et al. 2013). Interestingly, we observed significant changes in the coordination between cell rounding and NEP in the cancer cell lines (Fig. 1A-I), which resulted in an impairment of centrosome-nucleus axis repositioning. We hypothesized that abnormal cell rounding could be responsible for the defects in centrosome positioning that we observed in U2-OS and MDA-MB-468 cell lines. However, interfering with cell rounding in RPE-1 cells, either by delaying it or accelerating it, did not disrupt accurate centrosome positioning. These observations rule out mitotic cell rounding as the main driver of centrosome positioning in early mitosis and reinforce the efficiency of centrosome-nuclear axis orientation in RPE-1 cells. In addition, they also suggest that the cues instructing centrosome positioning emanate from the nucleus. Using the nucleus as a positional cue during early mitosis has significant advantages. This would allow centrosomes to orient correctly under conditions where external cues are mostly absent, due to the disassembly of adhesion complexes in late G2 (Dao et al. 2009; Matthews et al. 2012) and before the cortical loading of the LGN- -NuMa complex, which occurs only in prometaphase (Kiyomitsu and Cheeseman 2012; Kotak et al. 2012). In addition, this configuration would also enable the efficient assembly of a spindle scaffold (Nunes et al. 2020), ensuring a faster capture of kinetochores and mitotic progression (Kaseda et al. 2012; Silkworth et al. 2012; Nunes et al. 2020).

Many nuclear components could serve as potential candidates to influence centrosome behaviour. The LINC complex, due to its localization bridging the nucleus and cytoplasm, is ideally placed to perform this function. Accordingly, it is involved in centrosome-nucleus tethering and nuclear migration in *C. elegans* (Fridolfsson and Starr 2010) and myotubes (Cadot et al. 2012; Wilson and Holzbaur 2012), and was also shown to affect multiple aspects of mitotic progression (Magidson et al. 2011; Booth et al. 2019; Stiff et al. 2020; Belaadi et al. 2022). However, the molecular mechanism behind LINC complex-mediated centrosome positioning during the early stages of mitosis remained unclear. One likely possibility is that, during late G2, the LINC complex could be directly involved in the recruitment of dynein to the NE. It is well established that during this stage, two parallel and independent nucleoporin-mediated pathways regulate dynein NE loading (Hu et al. 2013). Upon phosphorylation by CDK1 (Baffet et al. 2015), the nucleoporin RanBP2 binds to BicD2, leading to dynein recruitment (Splinter et al. 2010; Gallisà-Suñé et al. 2023). Additionally, Nup133 recruits CENP-F, which then binds NudE/NudEL to load dynein on the NE (Bolhy et al. 2011). Once at the NE, dynein generates pulling forces on microtubules which results in centrosomes tethering to the nucleus, aiding centrosome separation (Raaijmakers et al. 2012) and facilitating NEP (Beaudouin et al. 2002; Salina et al. 2002). Here, we show that in untransformed RPE-1 cells, the LINC complex acts during late G2 to assist in NE- dynein loading (Fig. 5) and provide spatial cues for centrosome positioning (Figs. 4 and 5), in parallel with the NPC-mediated pathways. These observations fit nicely with a previous report showing a role for the LINC complex in centrosome movement during early mitosis (Stiff et al. 2020). In addition, LINC complex components SUN1 and SUN2, which are located on the inner nuclear membrane, were also shown to interfere with spindle assembly by delaying the removal of membranes from chromatin (Turgay et al. 2014; Belaadi et al. 2022). Together with our results, these observations indicate that the LINC complex acts on multiple levels during early spindle assembly. By assisting in the recruitment of NE-dynein, it ensures the accurate positioning of centrosomes on the shortest nuclear axis, to allow efficient assembly of the mitotic spindle. In line with this hypothesis, we and others have shown that disruption of the LINC complex, while unable to block mitotic entry, induces a severe delay in this process by decreasing the rate of cyclin B1 nuclear accumulation (Dantas et al. 2022) and interfering with spindle assembly efficiency (Turgay et al. 2014; Booth et al. 2019; Stiff et al. 2020). In parallel, by facilitating the removal of membranes from chromatin (Turgay et al. 2014), it also accelerates the capture of kinetochores by microtubules. Overall, this would result in a more efficient “search and capture” of chromosomes during early prometaphase which, together with the "ring-like” distribution of chromosomes (Magidson et al. 2011), ensures a maximum exposure of kinetochores to microtubules and decreases the probability of generating of erroneous kinetochore- microtubule attachments (Cimini et al. 2003; Silkworth and Cimini 2012). Notably, mutations and abnormal expression of LINC complex proteins have been implicated in a plethora of cancers (Sjöblom et al. 2006; Doherty et al. 2010; Matsumoto et al. 2015), suggesting a potential role in the maintenance of chromosomal stability. However, additional work is required to determine the exact molecular nature of the spatial cues that drive centrosome movement to the shortest nuclear axis. One hypothesis is that asymmetric localization of force-generating complexes on the NE could bias centrosome movement to a specific nuclear orientation. Accordingly, we observed an accumulation of SUN proteins in nuclear areas between the separating centrosomes, in a microtubule-dependent manner (Fig. 4). One alternative hypothesis is that centrosomes generate pushing forces on the nucleus, sufficient to deform it and create a shortest axis. If so, then the LINC complex would be necessary to ensure the timely loading of dynein on the NE, so that centrosomes could tether to the nucleus and drive its deformation. Indeed, centrosome-mediated NE invaginations were previously reported to occur during the G2-M transition (Georgatos et al. 1997; Beaudouin et al. 2002; Salina et al. 2002; Turgay et al. 2014) and help in NEP. Overall, we propose a model (Fig. 6) in which LINC complex-mediated loading of dynein at the NE dictates centrosome positioning at the shortest nuclear axis upon NEP and this is required to ensure an efficient spindle assembly in human cells.

## Supporting information

Reply to reviewers

## Acknowledgments

This work was funded by Portuguese funds through FCT—Fundação para a Ciência e a Tecnologia/Ministério da Ciência, Tecnologia e Ensino Superior in the framework of the project PTDC/BIA-CEL/6740/2020. J.T.L. is supported by grant SFRH/BD/147169/2019 from FCT. The authors thank Helder Maiato for critical reading of the manuscript and access to essential equipment for live cell imaging. Work in the Maiato lab is funded by the European Research Council consolidator grant CODECHECK, under the European Union’s Horizon 2020 research and innovation programme (grant agreement 681443), Fundação para a Ciência e a Tecnologia of Portugal (PTDC/MED-ONC/3479/2020) and La Caixa Health Research Grant (LCF/PR/HR21/52410025). The authors would like to thank Tom Misteli for the Halo-Tag9-LaminB1 plasmid, Jean de Gunzburg for the pRK5- Rap1[Q63E] construct and Stephen Royle for the plasmid pWPT LBR-mCherry.

## Author contributions

J.T.L. performed experimental work, analysed data, prepared figures and wrote the manuscript. A.J.P. developed the computational tool to measure fluorescence intensity at the NE and wrote the manuscript. J.G.F. provided the conceptual framework, prepared figures, wrote the manuscript and provided funding.

## Materials and Methods

### Cell lines

RPE-1, U2-OS and MDA-MB-468 cell lines were cultured in DMEM (Life Technologies) supplemented with 10% FBS (Fetal Bovine Serum; Life Technologies) and kept in culture in a 37°C humidified incubator with 5% CO_2_. For MDA-MB-468 cells, media was also supplemented with GlutaMAX^TM^ (Life Technologies). RPE parental, RPE H2B- GFP/mRFP-α-tubulin, U2OS parental, U2OS H2B-GFP/mRFP- α-tubulin and MDA-MB- 468 parental were already available in our lab. RPE-1 GFP-α-tubulin, RPE-1 GFP-α- tubulin/LBR-mCherry and MDA-MB-468 H2B-GFP/mRFP-α-tubulin cell lines were generated by transduction with lentiviral vectors containing the respective plasmids available in our lab – Table 1. For this purpose, we used HEK293T cells at a 50-70% confluence that were co-transfected with lentiviral packaging vectors (16.6μg of Pax2, 5.6μg of pMD2 and 22.3μg of the plasmid of interest), using 30μL of Lipofectamin 2000 (Life Technologies). Approximately 48 hours after transfection, the virus-containing supernatant was collected, filtered, and stored at -80°C. Cells were infected with the collected virus together with polybrene (1:1000) in standard culture media for 24h. Some days after the infection, the cells expressing the fluorescent tags were isolated by fluorescence activated cell sorting (FACS; FACS Aria II).

RPE-1 H2B-GFP/mRFP-α-tubulin/shSUN1 and RPE-1 H2B-GFP/mRFP-α- tubulin/shSUN2 were also generated by infecting cells, using commercially available lentiviral vectors encoding the desired shRNAs (MISSION® shRNA, Sigma-Aldrich; shSUN1 - TRCN0000133901 - Target Sequence: CAGATACACTGCATCATCTTT; shSUN2 - TRCN0000141958 - Target Sequence: GCAAGACTCAGAAGACCTCT). Cells were then selected with puromycin (20μg/mL; Merck Millipore) for 7 days. RPE-1 cells expressing the KASH constructs were generated via lentiviral infection. RPE-1 GFP-α-tubulin cells were infected with viruses containing the mCherry-DN-KASH or mCherry-DN-KASHΔPPPL plasmids, as described above. Doxycycline (5μg/ml; Fischer Scientific) was added for 24h to the culture media to stimulate the expression of the different KASH fusion proteins. After this period, the cells expressing the fluorescent tags were isolated by FACS and placed back in normal media for expansion. Addition of doxycycline was performed again 24 hours before imaging. RPE-1 H2B-GFP/mRFP-α- tubulin/Halo-Tag9-LaminB1 cell line was generated by transiently transfecting a HaloTag9-LaminB1 plasmid (gift from Tom Misteli) using Lipofectamin 2000 (Life Technologies). Specifically, 5μL of Lipofectamin 2000 and 0.6μg of HaloTag9-LaminB1 plasmid were diluted separated and incubated in OPTIMEM (Alfagene) for 30 min. The mixture was then added to confluent cells cultured and incubated for 6h in reduced serum medium (DMEM with 5% FBS). Cells were selected using 400μg/mL of G418 (Gibco, Life Technologies) for 14 days.

### Drug treatments

To visualise DNA, cells were incubated for at least 1 hour with SiR-DNA (Spirochrome), at a final concentration of 10nM, for a maximum of 4 hours of imaging. To visualise HaloTag9 fluorescence, Janelia Fluor (JF) Ligand-647 (Promega) was added to the imaging media at a final concentration of 75nM. To affect cell rounding, a ROCK inhibitor (Y-27632) was used at 20μM (Sigma-Aldrich) and cells were incubated with the drug 30-60 minutes prior to imaging. Calyculin-A (CalA, Abcam) was added to the cells during the imaging, at 20μM for RPE-1 cells and 50μM for U2-OS. To perturb microtubules, nocodazole (Noc, Sigma-Aldrich) was used at 3.3μM. Control cells were treated with the corresponding volume of DMSO (Sigma-Aldrich).

### Transfections

Cells were transfected with the plasmid pRK5-Rap1[Q63E] plasmid (Rap1*; a gift from Jean de Gunzburg) for 48 hours as described before for plasmid transfections. Control cells were transfected with Lipofectamin 2000 (Invitrogen) only, in the same conditions. To deplete LaminA from RPE-1 cells were transfected with small interfering RNAs (siRNAs) using Lipofectamin RNAi Max (Life Technologies). Specifically, 5μL of Lipofectamin RNAi Max and 20nM of each siRNA were diluted separately, and then incubated in OPTIMEM (Alfagene) for 30 min. The mixture was then added to confluent cells cultured and incubated for 6h in reduced serum medium (DMEM with 5% FBS). Commercial ON-TARGETplus SMARTpool siRNAis (Dharmacon) were used. Commercial ON-TARGETplus SMARTpool nontargeting siRNAs and mock transfections were used as controls. Cells were analysed 72h after transfection and protein depletion efficiency verified by immunoblotting.

### Western Blotting

Cell extracts were collected after trypsinisation and centrifuged at 1200rpm for 5min, washed in Phosphate Buffered Saline (PBS) twice and resuspended in 30-50μL of lysis buffer (50mM Tris-HCl pH7.4, 150mM NaCl, 1mM EGTA, 0.5% NP-40, 0.1% Triton X-100), 1:50 protease inhibitor (Complete Tablets EASYpack; 04693116001; Roche), 1:100 Phenylmethylsulfonyl fluoride (PMSF). Cells were kept on ice for 30 minutes, then flash frozen in liquid nitrogen twice. DNA was pelleted after centrifugation at 14,000 rpm for 8min at 4°C, the supernatant was collected, and protein concentration determined using the Bradford protein assay (Bio-Rad). Proteins were run on a 10% SDS-PAGE gel (30μg/lane) and transferred using a semi-dry blotting system (Trans-Blot Turbo system, Bio-Rad) for 10 minutes at 25V, with constant amperage. Next, the membranes were blocked with 5% milk in PBS with 0.05% Tween-20 (PBS-T) for 1h, at room temperature (RT). The primary antibodies used were anti-LaminA (C-terminal; 1:1000; L1293; Sigma-Aldrich), anti-SUN1 (1:500; MABT892; Merck Millipore), anti-SUN2 (1:500; MABT880; Merck Millipore), anti-GAPDH (1:20000; 60004-1-Ig; Proteintech), anti-β-tubulin (1:5000; Ab6046; Abcam). All primary antibodies were incubated overnight at 4°C with shaking. After three washes in PBS-T, the membranes were incubated with the secondary antibody for 1h, at RT. The secondary antibodies used were anti-mouse-HRP and anti-rabbit-HRP, at 1:5000. After three washes with PBS-T, the detection was performed with Clarity Western ECL Substrate (Bio-Rad). Acquisition of blots was performed with a Bio-Rad ChemiDoc XRS system using the IMAGE LAB software.

### Immunofluorescence

Cells were seeded the day before the experiment in coverslips coated with fibronectin (25 μg/ml; F1141; Sigma-Aldrich; FBN). When necessary, cells were treated with the appropriate compounds, and fixed with 4% paraformaldehyde (PFA) in cytoskeleton buffer (1.25M NaCl, 1M KCl, 125mM Na2HPO4, 250mM KH2PO4, 250mM EGTA, 250mM MgCl2, 250mM PIPES, 500mM Glucose, pH 6.1) for 10 minutes at RT, and then extracted in PBS with 0.5% Triton-X100 (Sigma-Aldrich) for 5 minutes (or 30 minutes when using the antibody against Dynactin 1). Coverslips were then blocked using 10% FBS in 0.1% Triton-X100 for 30 minutes, at RT. These coverslips were afterwards incubated with the following primary antibodies: rabbit anti-SUN1 (1:200; HPA008346; Sigma-Aldrich), rabbit anti-SUN2 (1:200; HPA001209; Sigma-Aldrich); mouse anti-Nesprin-2 (1:200; sc-398616; Santa Cruz Biotechnology), mouse α-tubulin (α-Tubulin B-5-1-2;1:1000; 32-2500; Sigma-Aldrich), rabbit β-tubulin (1:1000; Ab6046; Abcam); mouse anti-Lamin A+C (1:500; Ab8984; Abcam), rabbit anti-Lamin B1 (1:1000; ab16048; Abcam), rabbit α-Dynactin 1 (1:200;PA5-21289; Invitrogen), rabbit anti- Nesprin1 (1:200; PA5-82666; Invitrogen), anti-LBR (1:500; HPA062236, Atlas antibodies) in blocking solution. Primary antibody incubation was usually performed for 1 hour at RT, except when probing cells with the anti-Nesprin-1, anti-Nesprin2, and anti-Dynactin 1 antibodies, in which case incubation was completed overnight at 4°C. Coverslips were washed with PBS-0.1%Tríton-X three times (5 minutes each) and incubated with the secondary antibodies (1:2000; Alexa-Fluor conjugated; Invitrogen), at RT for 1h. When appropriate, DAPI (1μg/mL; Invitrogen) was added to the secondary antibody mixture to stain DNA. Finally, coverslips were washed three times in PBS with 0.1% Triton-X100 and once in PBS, and sealed on a glass slide using mounting medium (20nM Tris pH 8, 0.5 N-propyl gallate, 90% glycerol). Images were acquired using an AxioImager Z1 (63x, Plan oil differential interface contract objective lens, 1.4NA; all from carl Zeiss) which is coupled to a CCD camera (ORCA-R2; Hamamatsu Photonics) using the Zen software (Carl Zeiss).

### Plasmids

| Plasmid | Type | Origin |
| --- | --- | --- |
| pLKO.1 H2B-GFP | Lentiviral | Addgene |
| pRRL-mRFP-α-Tubulin | Lentiviral | In house |
| Rap1* | Transient Expression | Gift from Jean de Gunzburg |
| α-Tubulin-GFP | Lentiviral | In house |
| LBR-mCherry | Lentiviral | Gift from Stephen Royle |
| HaloTag9-LaminB1 | Transient Expression | Gift from Tom Misteli |
| pLKO.1 shSUN1 - puro | Lentiviral | Sigma-Aldrich |
| pLKO.1 shSUN2 - puro | Lentiviral | Sigma-Aldrich |
| pinducer 20 DN-KASH | Lentiviral | Addgene #125554 |
| pinducer 20 DN-KASHΔPPPL | Lentiviral | Addgene # 129280 |

### Micropatterning

Micropatterning was performed using a deep UV-light technique to normalise cell shape and adhesion area, as previously described(Azioune et al. 2009). Glass coverslips (square 22×22mm, 1.5, VWR; or round 25mm, 1.5, ThermoFisher) were activated with plasma (Zepto Plasma System, Diener Electronic) for 2 minutes. Following plasma treatment, coverslips were incubated with 0.2mg/mL PLL(20)-g[3,5]-PEG(2) (SuSoS) in 10 mM HEPES at pH 7.4, for 1 h, at RT. Coverslips were washed three times with water, and left to dry before being placed on a synthetic quartz photomask (Delta Mask), previously activated with deep-UV light (PSD-UV, Novascan Technologies) for 5 min, using a 3uL of MilliQ water to seal it to the mask. The coverslips were then irradiated through the photomask with the UV lamp for 5 min and left to dry before being incubated with FBN (25 μg/ml; F1141; Sigma-Aldrich), in 100 mM NaHCO3 at pH 8.6, for 30 minutes, at RT. Whenever possible, 5 μg/ml Alexa 647–conjugated fibrinogen (ThermoFisher Scientific) was added to the fibronectin mix in order to visualise the pattern surfaces. Cells were added to the freshly incubated coverslips and allowed to spread for 15 minutes, before removing excess cells and new culture medium added, and cells left to fully adhere for another 12-16 hours.

### Time lapse microscopy

Cells seeded in patterned or non-patterned surfaces are placed in Leibovitźs-L15 medium (Life Technologies), supplemented with 10% FBS and Antibiotic-Antimycotic Solution 100X (AAS; Life Technologies) right before imaging, alongside the corresponding drugs, where indicated. Live cell imaging experiments were performed using temperature-controlled Nikon TE2000 microscopes equipped with a modified Yokogawa CSU-X1 spinning-disk head (Yokogawa Electric), an electro multiplying iXon+ DU-897 EM-CCD camera (Andor) and a filter wheel. Three laser lines were used to excite 488nm, 561nm and 647nm and all the experiments were done with immersion oil using a 60x 1.4NA Plan-Apo DIC (Nikon). Image acquisition was controlled by NIS Elements Ar software. Images were obtained with 17 z-stacks (0.5μm step) with a 20 second interval when assessing centrosome positioning during mitotic entry.

### MATLAB custom algorithm for centrosome tracking

Analysis of centrosome positioning and behaviour during mitotic entry was performed using a custom-designed MATLAB (The MathWorks Inc, USA; v2018b) script(Castro et al. 2020) that contains a specialised workflow previously optimised for centrosome tracking, together with the reconstruction of both cellular and nuclear membranes in a 3D space. A pixel size of 0.176μm and a z-step of 0.5μm, were taken into consideration. Error correction methods were employed in cases where the standard automatic method was unable to correctly detect the two centrosomes and membranes. These correction methods involved, amongst others, manual centrosome position adjustment in all 3 coordinates (x, y and z) for each frame, and threshold correction for membrane reconstruction. From these membrane reconstructions, the algorithm was able to extract cell and nuclear major axis, as well as cell and nuclear membranes eccentricity and irregularity levels. From the coordinates obtained for centrosomes and with the nuclear and cell axis computed, the tool was able to calculate the angle between the two centrosomes that passed through the centroid of the nucleus (angle centrosome- centrosome), the angle of the centrosomes axis relative to the long nuclear axis (angle centrosome-nucleus), as well as the angle formed between the longest cell axis and the longest nuclear axis, for each frame/timepoint.

### Quantification of nuclear solidity

Nuclear solidity was quantified using the shape descriptor plugin from ImageJ. Briefly, nuclei are outlined using the polygon tool and nuclear area is measured. To calculate nuclear solidity, the nuclear area is then divided by the corresponding convex hull area. Irregular nuclei will typically show a lower nuclear solidity value.

### Quantitative image analysis of dynactin 1 and LINC complex proteins levels

For the quantification of dynactin 1, nesprin and SUN protein levels at the NE, images were analysed using ImageJ. A sum projection of three z-slices encompassing the central region of the nucleus was employed in all measurements. On the sum-projected image, a segmented line (smoothened by a spline fit) of a defined width (w1) was drawn along the NE, and the transverse-averaged fluorescence signal (S1), which contains signal as well as background, was measured. The S1 (transverse average) is directly retrieved using Ctrl-M or Ctrl-K in ImageJ. A second equivalent measurement (S2) was done using the same line after increasing its width to w2. While the signal of interest remains the same in the dilated line, background increases by the factor w2/w1, the knowledge of which allows retrieval of I(r), the background-corrected profile, via

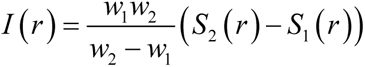

Line width w1 should be large enough to fully encompass the signal of interest, while w2 should be at least 20% larger than w1 but small enough to avoid inclusion of extraneous signal from non-NE sources. In the particular quantification done in this study, the intensity profile (i.e. the r-dependence) was irrelevant, so I(r) was integrated along the full length of the curve and divided by the line length (or, equivalently, the line ’area’).

### Statistical analysis

At least three independent experiments were used for statistical analysis. Sample sizes and number or replicates are indicated in each figure legend. Normality of the samples was assessed using the Kolmogorov–Smirnov test. Statistical analysis for multiple group comparison was performed using a parametric one-way analysis of variance (ANOVA) when the samples had a normal distribution. Otherwise, multiple group comparison was done using a nonparametric ANOVA (Kruskal-Wallis). Multiple comparisons were analyzed using either post-hoc Student-Newman-Keuls (parametric) or Dunn’s (nonparametric) tests. When comparing two experimental groups only, a parametric *t* test was used when the sample had a normal distribution, or a nonparametric Mann–Whitney test was used for samples without normal distribution. Comparison for multiple time-course datasets was done using an ANOVA Repeated Measures, when the samples had a normal distribution. Otherwise, group comparison was done using Repeated Measures ANOVA on Ranks. No power calculations were used. All statistical analyses were performed using the GraphPad Prism (Dotmatics). When comparing proportions between 2 populations, a z-score was calculated.

---

**Supplementary Figure S1 –Workflow and output derived from the custom-designed MATLAB script (Trackosome)**

**(A)** Layout of the interface utilised to automatically track centrosomes, and its various manual correction options - “Correct Centrosome Coordinates”. Each rectangle of the progress bar, shown on the lower section of the image, represents a time point in the movie analysed (blue – automatically tracked, grey – manually corrected, yellow – one or both centrosomes slightly out of focus), for this RPE-1 cell (same shown in Figure 1A). Centrosome 1 represented in blue, and centrosome 2 represented in red. **(B)** Representative graph showing the location of each centrosome in time and 3D space (x, y and z). **(C)** Reconstruction of cell membrane (green) and nuclear membrane (yellow) in 3D, as well as centrosome location relative to these two structures (blue and red dots) obtained for one of the time frames of the movie. **(D)** Representative graph displaying the variation in the “angle centrosome-centrosome” over time. This angle is calculated by a vector that links the two centrosomes and intercepts the centroid of the nucleus. **(E)** Representative graph displaying the variation in the “angle cell-nucleus” (alignment of the long cell axis and long nuclear axis; red), “angle centrosome-cell” (alignment of the centrosomes vector with the long cell axis; green) and “angle centrosome-nucleus” (alignment of the centrosomes vector with the long nuclear axis; blue), over time. **(F)** Scheme depicting how the alignment between longest cell axis and longest nuclear axis is calculated. **(G)** Scheme depicting how the alignment between the centrosome pair and the longest nuclear axis is calculated. **(H)** Scheme depicting how centrosome separation is calculated. All calculations are performed using datasets previously processed with Trackosome. **(I)** Representative polar plot that displays the distribution of centrosomes positioning relative to the longest nuclear axis at the moment of NEP. Note that an orientation towards 90° corresponds to alignment with the shortest nuclear axis. **(J)** Representative plot displaying the correlation between centrosome separation (angle centrosome-centrosome) and positioning on the shortest nuclear axis (angle centrosome-nucleus) at the moment of NEP. Note that when centrosomes are on opposite sides of the nucleus and on the shortest nuclear axis, data points will cluster on the top right corner of the graph (green box). When centrosomes do not separate, data points will cluster on the bottom left corner (red box).

**Supplementary Figure S2 – External cues do not impact centrosome positioning on the shortest nuclear axis at NEP**

**(A)** Representative frames of the moment of NEP from movies of Mock or Rap1* transfected MDA-MB-468 cells, stably expressing H2B-GFP and RFP-tubulin, seeded on a 10μm wide-micropatterned line. **(B)** Quantification of cell membrane eccentricity of Mock transfected cells (gray; n= 25) and Rap1* expressing cells (pink; n= 9; ****p<0.001). Dashed line represents moment of NEP. Correlation between the angle centrosome-centrosome (x axis) and the angle centrosome-nucleus (y axis) at the moment of NEP for Mock **(C)** and Rap1* **(D)** transfected cells. Representative frame of the moment of NEP from U2-OS cells, stably expressing H2B-GFP and RFP-tubulin, seeded on a 10μm wide-micropatterned line, treated with DMSO **(E)**, Y-27632 **(F)**, CalyculinA **(G**; CalA). Quantification of cell membrane eccentricity for DMSO (**H**; gray; n=32) and Y-27632 treated cells (green; n=32; ****p<0.001) or DMSO (**I**; gray) and CalA treated U2-OS cells (green; n=22; ****p<0.0001). Polar plots quantifying centrosome positioning relative to the longest nuclear axis at NEP for U2-OS cells treated with DMSO **(J)**, Y-27632 (**K**; p=0.683) and CalA (**L**; p=0.282). **(M)** Correlation between the angle centrosome-centrosome (x axis) and the angle centrosome-nucleus (y axis) at the moment of NEP, for cells treated with DMSO (top graph), Y-27632 (middle graph) and CalA-treated (bottom graph). Representative frame of the moment of NEP from U2-OS cells, stably expressing H2B-GFP and RFP-tubulin seeded on a 20μm wide- micropatterned line **(N)** or a non-patterned, fibronectin (FBN)-coated surface **(O)**. Correlation between the angle centrosome-centrosome and the angle centrosome-nucleus for cells seeded on 20μm wide-lines (**P**; n=32) or non-patterned FBN-coated surface (**Q**; n=34). Polar plots quantifying centrosome positioning relative to the longest nuclear axis at NEP for the cells seeded on 20μm wide-lines (**R**) or non-patterned, FBN-coated surface (**S**). **(T)** Quantification of cell membrane eccentricity of DMSO treated cells (gray), cells seeded on 20μm wide-lines (black; ****p<0.001) and non-patterned, FBN-coated surface (green; ****p<0.001). Scale bars for all images, 10μm.

**Supplementary Figure S3 – Nuclear lamina levels are altered in CIN cells**

Representative images of RPE-1 (top panel), U2-OS (middle panel) and MDA-MB-468 (bottom panel) cells, immunostained for Lamin A/C **(A)** and LaminB1 **(B)**. **(C)** Quantification of NE fluorescence intensity of LaminA/C of RPE-1 (n= 60), U2-OS (n= 67; ****p<0.001) and MDA-MB-468 cells (n= 50; n.s. – not significant). **(D)** Quantification of the NE fluorescence intensity of Lamin B1 of RPE-1 (n= 50), U2-OS (n= 50; ****p<0.001) and MDA-MB-468 cells (n= 51; ****p<0.001). **(E)** Quantification of the ratio of NE intensity of Lamin A/C:Lamin B1 for each of the cell lines, calculated from the values obtained in (C) and (D). **(F)** Quantification of the Lamin A/C:Lamin B1 ratio calculated from expression levels obtained from the online repository DepMap.com. **(G)** Quantification of nuclear solidity levels for RPE-1, U2-OS (****p<0.001) and MDA-MB- 468 (****p<0.001) cells used in (C). Nuclear solidity was measured in Fiji and is defined as nucleus area/nucleus convex area. **(H)** Quantification of Lamin B1 (LMNB1) levels in control and LMNB1 overexpressing cells, using immunofluorescence (****p<0.001). **(I)** Quantification of LBR expression levels in control and LBR-overexpressing cells, using immunofluorescence (****p<0.001).

**Supplementary Figure S4 - Depletion of SUN1 and SUN2 using shRNA generates a heterogenous population of cells**

**(A)** Representative images of Control and shSUN1-treated cells, stably expressing H2B-GFP and RFP-tubulin seeded on FBN and immunostained for SUN1. **(B)** Western blot to confirm the efficiency of SUN1 depletion in the overall population. **(C)** Representative images of Control and shSUN2-treated cells, stably expressing H2B- GFP and RFP-tubulin seeded on FBN, immunostained for SUN2. **(D)** Western blot to confirm the efficiency of SUN2 depletion in the overall population. Yellow arrows indicate cells with higher depletion levels. **(E)** Expression of the DPPPL-KASH (top panel) and DN-KASH (bottom panel) constructs in RPE-1 prophase cells. Yellow arrows indicate the NE. **(F)** Representative image of a control RPE-1 cell in prophase, immunostained for Nesprin 1 and Nesprin 2, highlighting their NE localization. For all images, scale bars represent 10mm. **(G)** Representative immunofluorescence images from DPPPL-KASH (top panel) and DN-KASH (bottom panel), to highlight LMNB1 localization. **(H)** Quantification of LMNB1 levels between DPPPL-KASH and DN-KASH (n.s. – not significant).

**Supplementary Figure S5 – Analysis of LINC complex components in U2-OS and MDA-MB-468 cell lines**

**(A)** Representative immunofluorescence images showing the NE localization of nesprin 1 (SYNE1) and nesprin 2 (SYNE2) for RPE-1 (top panel), U2-OS (middle panel) and MDA-MB-468 (bottom panel) cell lines. Quantification of the levels of SYNE1 **(B)**, SYNE2 **(C)**, SUN1 **(D)** and SUN2 **(E)** obtained from immunofluorescence images for all cell lines (****p<0.001; **p<0.01; n.s. – not significant). **(F)** Representative immunofluorescence images showing the NE localization of dynactin 1 in RPE-1 (top panel), U2-OS (middle panel) and MDA-MB-468 (bottom panel). **(G)** Quantification of the levels of dynactin 1 on the NE obtained from immunofluorescence images for all cell lines (****p<0.0001). Scale bars for all images, 10 mm.

**Supplementary Movie S1 – Mitotic entry of an RPE-1 seeded on a line micropattern**

RPE-1 cell expressing histone H2B-GFP (magenta) and tubulin-RFP (gray) seeded on a 10mm wide line fibronectin (FBN) micropattern, showing top and lateral projections. Time lapse is 20sec.Time is in min:sec and time zero corresponds to NEP. Scale bar=10mm.

**Supplementary Movie S2 – Mitotic entry of an U2-OS cell seeded on a line micropattern**

U2-OS cell expressing histone H2B-GFP (magenta) and tubulin-RFP (gray) seeded on a 10mm wide line fibronectin (FBN) micropattern, showing top and lateral projections. Time lapse is 20sec. Time is in min:sec and time zero corresponds to NEP. Scale bar=10mm.

**Supplementary Movie S3 – Mitotic entry of a MDA-MB-468 cell seeded on a line micropattern**

MDA-MB-468 cell expressing histone H2B-GFP (magenta) and tubulin-RFP (gray) seeded on a 10mm wide line fibronectin (FBN) micropattern, showing top and lateral projections. Time lapse is 20sec. Time is in min:sec and time zero corresponds to NEP. Scale bar=10mm.

**Supplementary Movie S4 – Mitotic entry of an RPE-1 seeded on a FBN-coated coverslip**

RPE-1 cell expressing histone H2B-GFP (magenta) and tubulin-RFP (gray) seeded on non-patterned (FBN) micropattern, showing top and lateral projections. Time lapse is 20sec. Time is in min:sec and time zero corresponds to NEP. Scale bar=10mm.

**Supplementary Movie S5 – Mitotic entry of an RPE-1 depleted of SUN1 seeded on a FBN-coated coverslip**

RPE-1 cell expressing histone H2B-GFP (magenta) and tubulin-RFP (gray) and treated with shSUN1, seeded on non-patterned (FBN) micropattern, showing top and lateral projections. Time lapse is 20sec. Time is in min:sec and time zero corresponds to NEP. Scale bar=10mm.

**Supplementary Movie S6 – Mitotic entry of an RPE-1 expressing DPPPL-KASH seeded on an FBN-coated coverslip**

RPE-1 cell expressing histone tubulin-GFP (gray), DPPPL-KASH-mCherry (cyan) and SiR-DNA (magenta) seeded on non-patterned (FBN) micropattern, showing top and lateral projections. Time lapse is 20sec. Time is in min:sec and time zero corresponds to NEP. Scale bar=10mm.

**Supplementary Movie S7 – Mitotic entry of an RPE-1 expressing DN-KASH seeded on an FBN-coated coverslip**

RPE-1 cell expressing histone tubulin-GFP (gray), DN-KASH-mCherry (cyan) and SiR- DNA (magenta) seeded on non-patterned (FBN) micropattern, showing top and lateral projections. Time lapse is 20sec. Time is in min:sec and time zero corresponds to NEP. Scale bar=10mm.

