## Supplementary figures and images for "The LINC complex ensures accurate centrosome positioning during prophase"

### Supplementary Figure 1

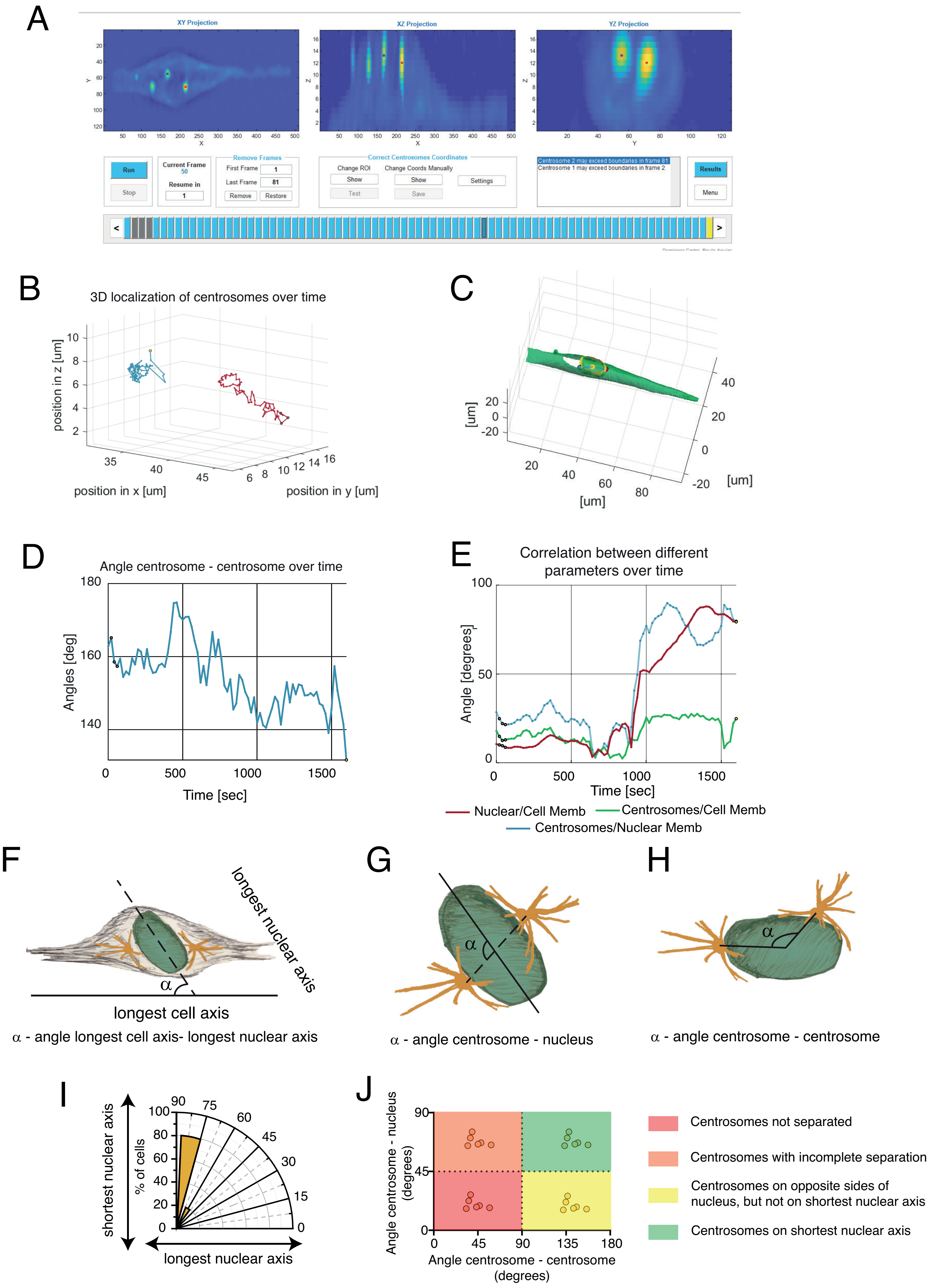


#### Supplementary Figure 1

### Supplementary Figure 2

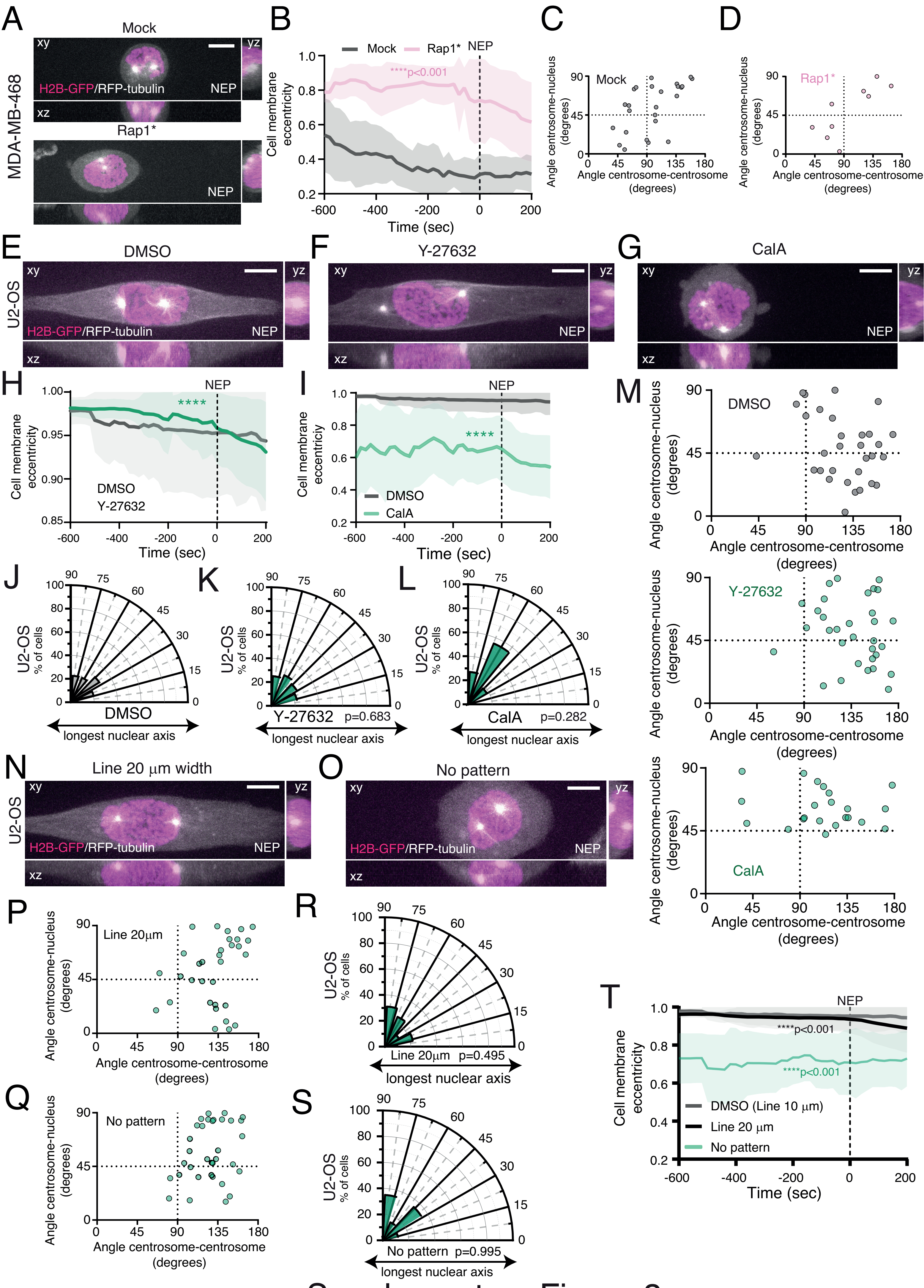


Supplementary Figure 2

### Supplementary Figure 3

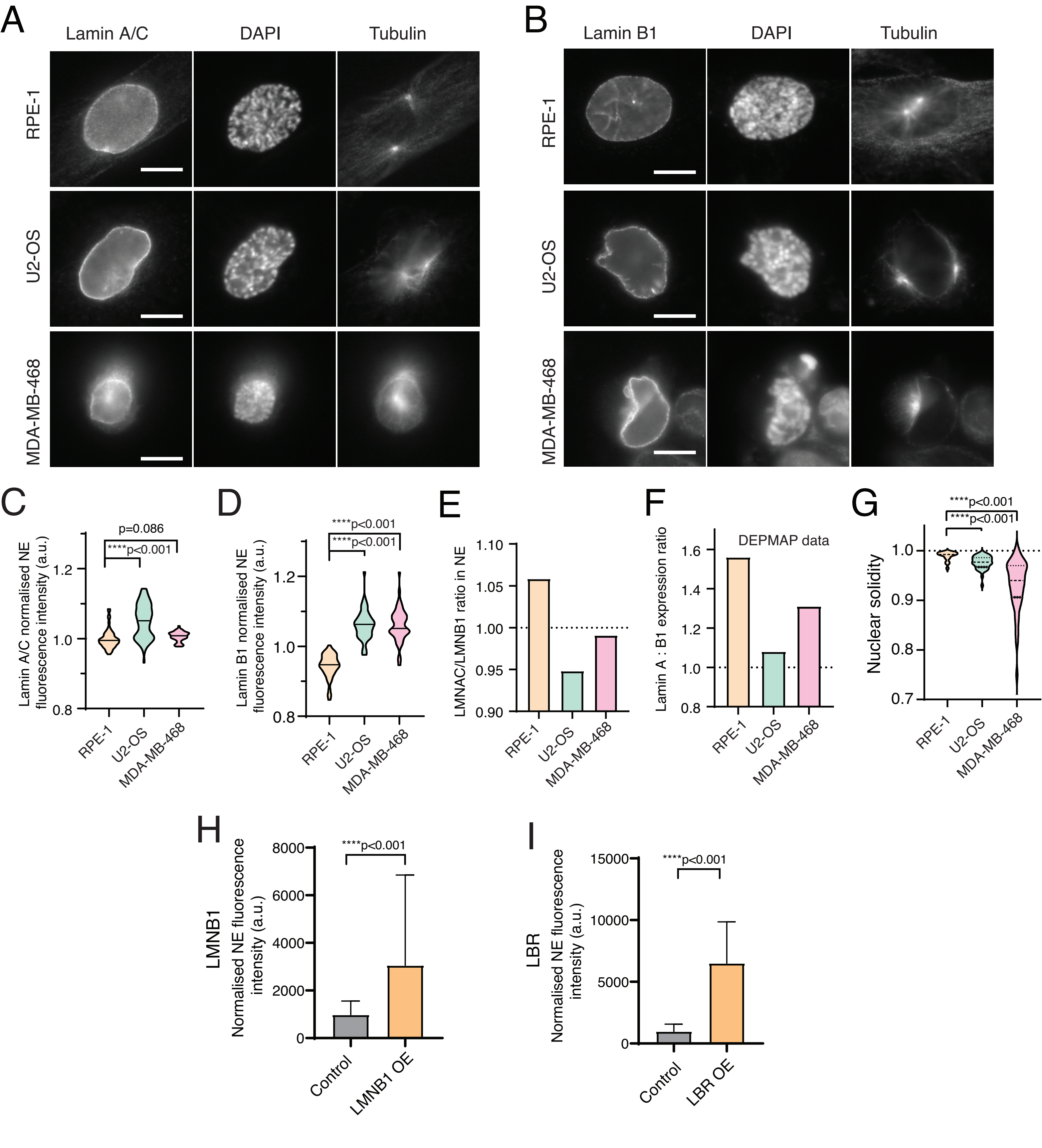


Supplementary Figure 3

### Supplementary Figure 4

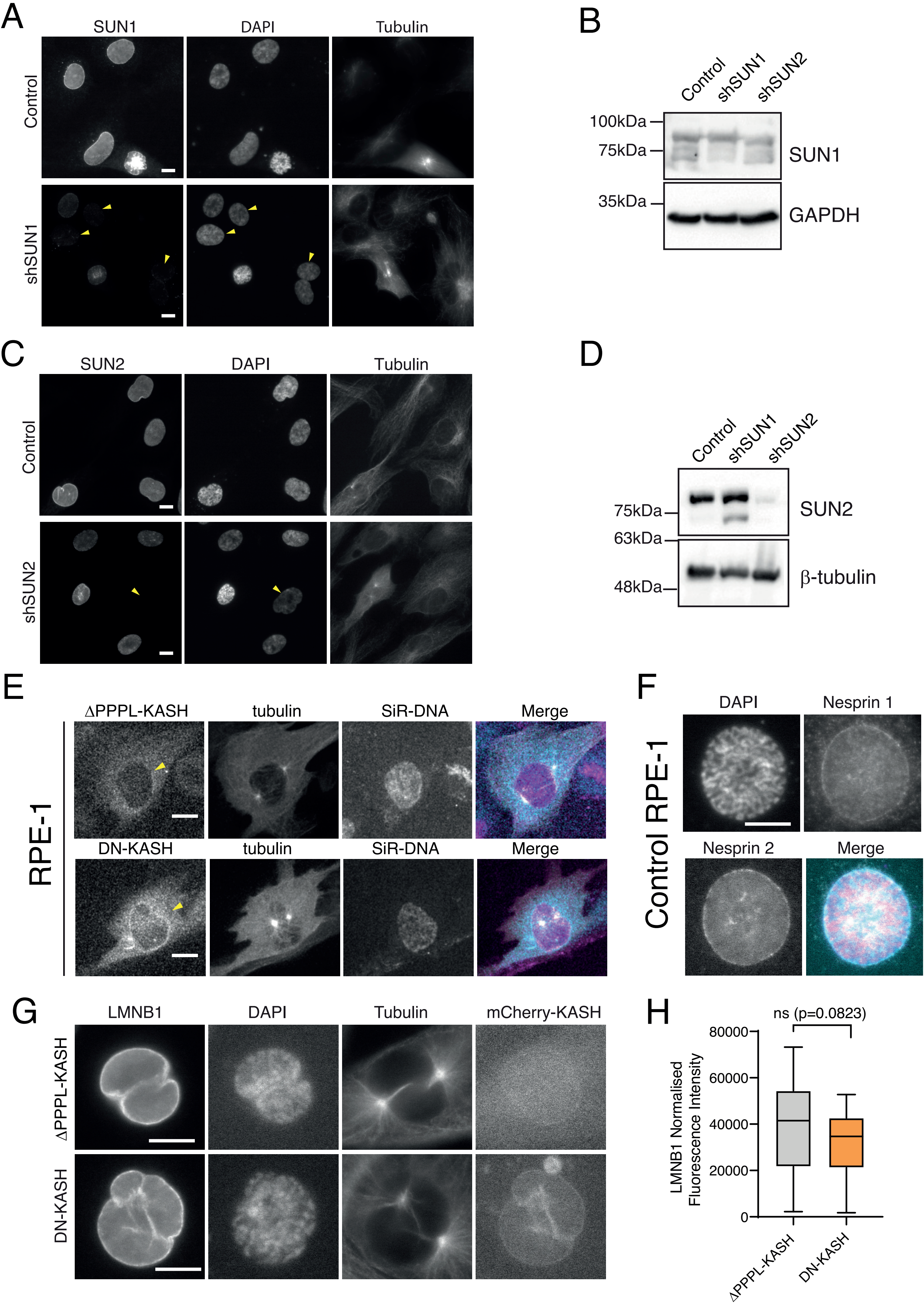


Supplementary Figure 4

### Supplementary Figure 5

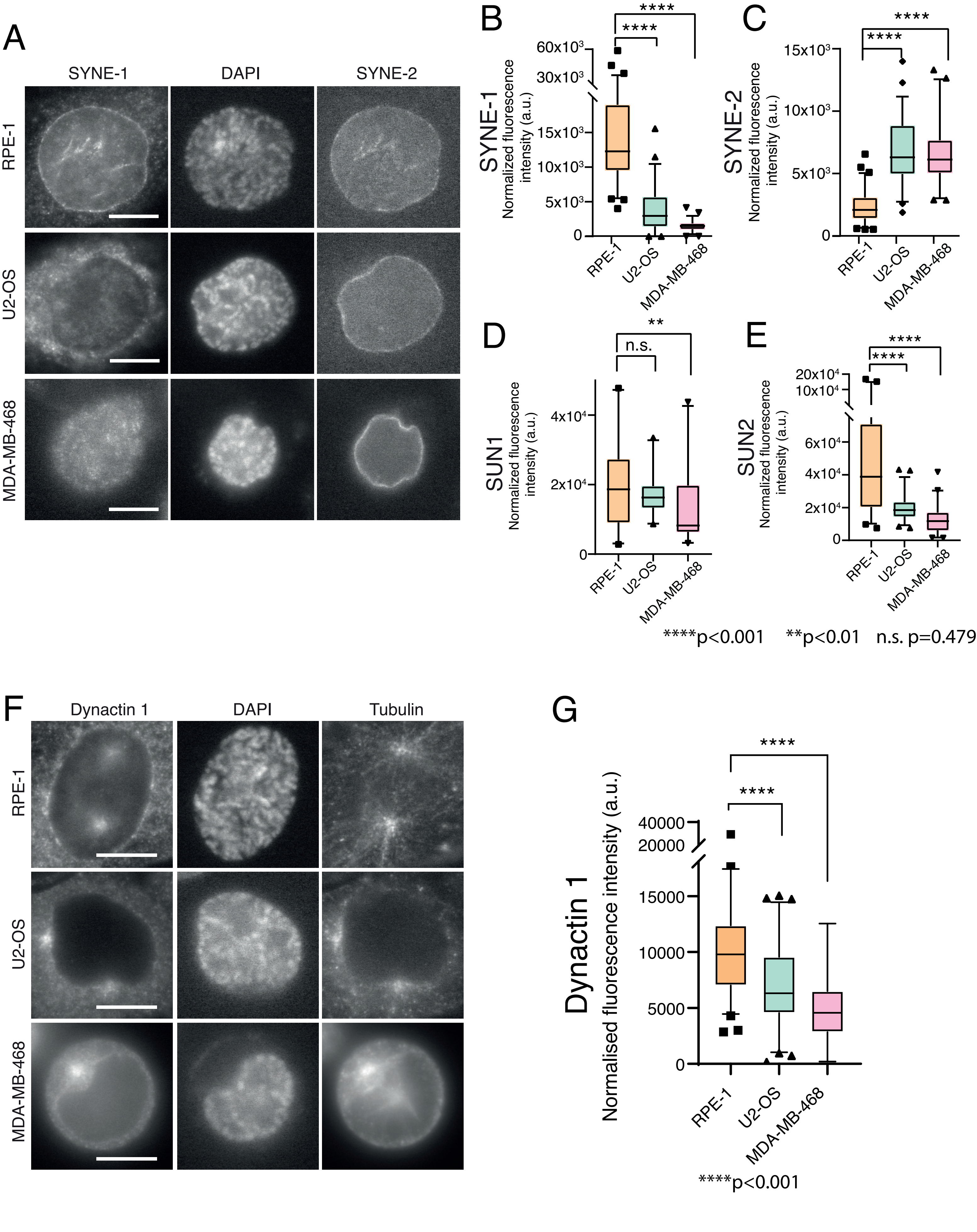


Supplementary Figure 5
