## Supplementary material for "The LINC complex ensures accurate centrosome positioning during prophase": Reply to reviewers

**Point-by-point reply to the reviewers` comments and suggestions**

**Reviewer 1:**

*- "Moreover, we demonstrate this mechanism is altered in cancer cells, leading to increased chromosome segregation errors. » Here the authors infer that the identified mechanism is absent in cancer cells and that its absence contributes to chromosome segregation errors. Both conclusions are not supported by the presented data. First, the authors did not test whether any members of the LINC complex or dynactin is present at lower levels on the nuclear membranes of the cancer cells. Such a direct validation would be essential to make such a strong statement. Second, the authors conclude that this mechanism prevents chromosome segregation errors, based on the fact that depletion or impairment of the LINC complex (shSUN1, shSUN2, DN-KASH) results in chromosome segregation errors. These perturbations lead, however, as noted by the authors themselves to pleiotropic effects, including insufficient retraction of nuclear membrane, which can all contribute to chromosome segregation errors. It is therefore impossible to estimate the contribution of the centrosome positioning mechanism to these segregation errors using this type of perturbations. One could even argue that this mechanism might not be that important, since depletion of SUN2, which also impairs centrosome positioning has no significant effect on chromosome segregation.*

We agree with the reviewer that an analysis of the levels of LINC complex components and dynactin in cancer cells is lacking. For this reason, we have now analyzed the levels of SUN1, SUN2, dynactin and nesprins by immunofluorescence in RPE-1 as well as the cancer cell lines. This was added to a new supplementary figure S5 and is discussed in the text.
In addition, considering the reviewer´s concern related to the pleiotropic effects of LINC complex depletion on chromosome segregation (with which we fully agree), we decided to remove this section and rewrite the manuscript and conclusions to focus on the role of the LINC complex in centrosome positioning only.

*- "The authors conclude based on three cell lines that the centrosome positioning mechanisms is present in non-transformed cells and not in cancerous cells. The authors have, however, only analysed 1 non-cancerous cell line, and they compare cells originating from vastly different tissues (retina, bones and breast) and origins (epithelial vs. mesenchymal cartilage cells). Such*

INSTITUTO
DE INVESTIGAÇÃO
E INOVAÇÃO
EM SAÚDE
UNIVERSIDADE
DO PORTO

Rua Alfredo Allen, 208
4200-135 Porto
Portugal
+351 220 408 800

www.i3s.up.pt

*a general statement is not possible without a systematic comparison of several healthy cells vs cancerous cells from the same tissue".*

We agree with this reviewer´s comment, which is also shared by the other reviewers. Accordingly, we have now extensively rewritten the manuscript to tone down this statement and focus on the role of the LINC complex in determining centrosome positioning.

*- "While the data showing that centrosome positioning depends on the LINC complex is solid and robust, some of the "negative" examples identified by the authors are less convincing. One the process the authors study is cell rounding. Based on the fact that Rap1 transfection or treatment with Calyculin A does not lead to differences that are statistically different, the authors conclude that cell rounding is not involved. However, absence of statistical difference does not mean that there is no difference. Indeed, when comparing the raw data in Figure 2L and 2Q to the positive hit shSun2 in Figure 4J, one could conclude that cell rounding does make a difference, and that this statistical difference would emerge if the authors would count a high number of cells. Therefore the authors should interpret these results in a more differentiated manner, and also instead of just stating nonsignificant, state also the real p-values for the different experiment".*

According to the reviewer´s suggestion, we have now added all p values to the respective graphs and interpreted these results in a more step-by-step manner. Moreover, while we understand the reviewer`s comment regarding our sample size, it should be noted that this is a single-cell, high-resolution imaging approach which, in combination with certain treatments makes it very challenging to obtain data for a high number of cells. In this regard, we point out that interfering with cell rounding was extremely difficult to achieve. When highly overexpressed, Rap1* completely impairs mitotic cell de-adhesion, and this blocks mitotic entry (Marchesi et al., 2014). Furthermore, CalA treatment induces a fast and drastic rounding, which makes it very challenging to accurately track centrosome and nuclear positions. Nevertheless, we filmed additional cells treated with CalA and added the data to the figures. Our results still confirm that interfering with cell rounding does not significantly change centrosome positioning during this stage. We would also like to note that the sample size in all conditions is within the range normally used when performing single-cell high resolution imaging.



INSTITUTO
DE INVESTIGAÇÃO
E INOVAÇÃO
EM SAÚDE
UNIVERSIDADE
DO PORTO

Rua Alfredo Allen, 208
4200-135 Porto
Portugal
+351 220 408 800

www.i3s.up.pt

*- The second major concerns emerges when looking at the data in Figure 5, when the authors test for the abundance of the dynein complex on the nuclear envelope in cells treated with DPPPL-KASH or DN-KASH. Yes, there is a statistical difference, but the absolute difference is tiny (I estimated a normalized intensity of 1.44 vs 1.35). This is a difference of less than 10%. How do the authors think that such a small change in dynein could have such a strong effect on centrosome positioning? Would a partial dynactin depletion by 10% give an equivalent result? Does the depletion of other proteins involved in the late recruitment of dynein at the NE also affect centrosome positioning?*

We thank the reviewer for this important point. Originally, we quantified dynactin intensity by selecting three unbiased random regions of the NE. However, it is possible this approach underestimates the overall fluorescence intensity across the entire structure. For this reason, we have now developed a custom-designed MATLAB algorithm to measure dynactin fluorescence intensity over the entire NE using the same dataset. We have replaced Fig. 5K and L with this data and a description of the method has been added to the Materials and Methods section. As can be seen from the new graph, there is a reduction of approximately 50% in dynactin NE fluorescence intensity.
The reviewer also asks whether depletion of other proteins involved in the late recruitment of dynein at the NE would also affect centrosome positioning. However, extensive previous work done by us and others, has shown that depletion of either BicD2 or NudE/NudEL, which are the main adaptors for dynein loading during the G2/M transition, significantly affect prophase centrosome positioning, since they detach centrosomes from the NE (Splinter et al., 2010; Bolhy et al., 2011; Hu et al., 2013; Baffet et al., 2015; Nunes et al., 2020). Once detached, centrosomes are no longer able to orient according to nuclear cues (Nunes et al., 2020). Therefore, we do not believe such an approach would provide additional information regarding the role of the LINC complex in this process.

**Reviewer 2**

*“The authors need some other NE protein as a control to show that the reduction of dynein by DN-KASH is a specific defect and not a broad impact on the NE. The dynein data in Figs. 5J-L need to be extended to SUN1/2”.*

We thank the reviewer for these suggestions. To clarify this point, we have analyzed the levels of lamin B following expression of DN-KASH or □PPPL-KASH. This data was added to Fig. S4. This allowed us to conclude that expression of the DN-KASH construct affects dynein, but likely does not impact

INSTITUTO
DE INVESTIGAÇÃO
E INOVAÇÃO
EM SAÚDE
UNIVERSIDADE
DO PORTO

Rua Alfredo Allen, 208
4200-135 Porto
Portugal
+351 220 408 800

www.i3s.up.pt

other NE proteins. In addition, we have also analyzed dynactin levels following SUN1 and SUN2 depletion. This data have been included in Figure 5.

*- "Figure 1 is insufficiently explained. The authors have to describe in an understandable way how they measured centrosome-centrosome angle and centrosome-nucleus angle. They should show a cartoon in which these angles are clearly shown. The small cartoons in Fig. 1C are not helpful at all; they are also not explained. The authors should explain the meaning of the black dots (are these centrosomes?) and the even smaller dots. The short nuclear axis should be indicated, e.g., by a red line".*

We apologize for the lack of sufficient explanation in Figure 1. We have now re-written the text to clarify all the points. We have also added a scheme explaining how centrosome-nucleus and centrosome-centrosome angles are quantified, according to the reviewer´s suggestion. We have added this to Fig. S1. We believe this makes our data more understandable and easier to follow.

*"On the first page of the manuscript: "Consequently, at the NEP, centrosomes are positioned on the shortest nuclear axis (Fig. 1C) as can be seen in Fig. 1A. This means that the centrosome-nucleus angle relative to the shortest nuclear axis should be 0. However, in Fig. 1C, this angle is between 45 and 90 degrees. This is also the case for Fig. 1D. Please clarify".*

We thank the reviewer for noticing this error. In fact, the graphs should always reflect positioning of centrosomes relative to the longest nuclear axis. Therefore, when the values are close to 90º, this means they are oriented on the shortest nuclear axis. We understand this could be confusing to the readers. We have now clarified this information throughout the text.

*- "I find it confusing that in Fig. 1, depending on the subfigure, the short or longest nuclear axis is used as a reference point: Fig. 1C: shortest; D: shortest; F: shortest; G: longest; I: shortest; J: longest. Thus, even within the same cell line, the reference point is changing. What is the rational for this variation"?*

Again, we refer to the point above. The reference point is always the shortest nuclear axis. However, we apologize for the lack of clarity. This has all been changed according to the explanation provided in the previous point.

*- Fig. 4K, L, M: in figure, y-axis: "shortest nuclear axis". In legend: "relative to the longest nuclear axis". I guess the longest nuclear axis is correct. Same in Fig. 5D and E. Fig. 5C lacks the WT control.*

INSTITUTO
DE INVESTIGAÇÃO
E INOVAÇÃO
EM SAÚDE
UNIVERSIDADE
DO PORTO

Rua Alfredo Allen, 208
4200-135 Porto
Portugal
+351 220 408 800

www.i3s.up.pt

This information has been clarified in the text and panels have been corrected accordingly. Regarding Fig. 5C, we believe the correct control is the expression of [illegible]PPPL-KASH, since it has been shown extensively that nesprins localize to the NE in control, unmanipulated cells. Nevertheless, we have added a WT control to Supplementary Figure 5, showing localization of nesprins in unmanipulated prophase cells.

*“The cells in Fig. 5J are not comparable: one has a monopolar spindle, the other a bipolar. The authors need some other NE protein as a control to show that the reduction of dynein by DN-KASH is a specific defect and not a broad impact on the NE. The dynein data in Figs. 5J-L need to be extended to SUN1/2”.*

We agree with the reviewer´s comment that the cell in the top panel might appear as a monopolar. However, it is not. In fact, this cell has centrosomes on the top and bottom of the nucleus, in a vertical configuration, a common feature of RPE-1 cells (check Magidson et al., Cell, 2011). To clarify this, we have now added lateral projections of all cells, highlighting the centrosomes to clearly show they are positioned on opposite sides of the nucleus.
The other points related to the effects of DN-KASH on other NE proteins and dynactin levels following shSUN1 and shSUN2 were also addressed and added to the figures.

*“The title of the paper is misleading: the authors do not provide any indication for a nuclear signal in prophase that determines centrosome positioning”.*

We have changed the title of the manuscript according to the reviewer´s suggestion.

*“It would make sense to use the same time scale in Figs. 1A and B (either min.sec. or sec.) to allow direct comparison”.*

We have now changed the time scale to seconds in all figures to allow direct comparison.

*“2nd section: Mitotic cell rounding "The authors state: Given that cancer cells failed... I would be careful with this generalization; only one cancer cell was used in this study”.*

Given the limited number of cells that we used, and following the concern raised by all reviewers, we have now re-written the text to avoid generalizations regarding cancer cell lines. Instead, we now focus on the role of the LINC complex in determining centrosome positioning.

INSTITUTO
DE INVESTIGAÇÃO
E INOVAÇÃO
EM SAÚDE
UNIVERSIDADE
DO PORTO

Rua Alfredo Allen, 208
4200-135 Porto
Portugal
+351 220 408 800

www.i3s.up.pt

A01/00

*“The authors say: "However, they did not place the centrosomes at the shortest nuclear axis (Figure 4K-M)." Centrosomes are still on the shortest nuclear axis but not as frequent as in control”.*

This has been corrected.

*“The white color in Fig. 6B cannot be seen and needs to be changed to something else”.*

We apologize for this oversight. During the upload and pdf conversion process, we did not realize the color of this bar, corresponding to the DN-KASH group had changed to white. However, given the concerns raised by reviewer 1 relative to the pleiotropic effects of SUN depletion, we have now removed the data concerning chromosome segregation and focus instead on the centrosome positioning mechanism.

*The paper has neither line nor page numbers.*

This has been added.

**Reviewer 3:**

*“Fig. 3: I suggest to quantify the lamin B1 and LBR overexpression levels”.*

According to the reviewer´s suggestion, we have quantified the levels of lamin B1 and LBR in cells overexpressing the respective constructs. These data were added to Fig. S3.

*“it would make sense to indicate the test used for each p-value in all the figure legends”.*

We have now added the statistical test used and the p-value in the figure legends.

*“Figure legends are quite repetitive and could be shortened. E.g. in Fig. 1 the description for E, F, H and I repeats what has been explained for B and C. Same*

INSTITUTO
DE INVESTIGAÇÃO
E INOVAÇÃO
EM SAÚDE
UNIVERSIDADE
DO PORTO

Rua Alfredo Allen, 208
4200-135 Porto
Portugal
+351 220 408 800

www.i3s.up.pt

*applies between figure legends. The authors might refer to previous legends if the analysis was done in a similar way”.*

We have now simplified the legends.

*“How is nuclear solidity defined and analyzed in Fig S3D”?*

Nuclear solidity was analyzed using Fiji. In short, nuclei are outlined using the polygon tool and nuclear area is measured. To calculate nuclear solidity, the nuclear area is then divided by the corresponding nuclear convex hull area. Irregular nuclei will typically show a lower nuclear solidity value. This information was added to the manuscript.

*“The references to Fig S3 in figure legend 3 ("see Fig S3") do not enlighten the message and could be removed. The same applies to Fig5 - here it is not clear why the author refer to Fig S4”.*

We agree with this reviewer´s comment. We have now removed these references from the legends.

*“Fig. 5: Consider reordering the panel: Start with the current panel C (as in the text) as it is the necessary control prior to the experimental data”.*

We have now changed the order of the panel according to the reviewer´s suggestion.

*“Fig 5 I: what means "before"? Can the authors give a time window they use for analysis”.*

We have now replaced the term “before” with a defined time.

*“Page 20: "... shortest nuclear axis (Fig. 1C, 5D-G; n.s. - not significant). However, DN-KASH-expressing cells showed compromised separation and positioning of centrosome (Fig. 5D-G, * p=0.0155 and * p=0.0237, respectively). - rather point to the specific panels, i.e. Fig. 1C, 5D and F as well as and Fig. 5E and G”.*

We have now clarified these points in the text.

INSTITUTO
DE INVESTIGAÇÃO
E INOVAÇÃO
EM SAÚDE
UNIVERSIDADE
DO PORTO

Rua Alfredo Allen, 208
4200-135 Porto
Portugal
+351 220 408 800

www.i3s.up.pt

*“Fig 6B. The DN- KASH bars are on my pdf not visible - use a darker grey”.*

As mentioned above, we apologize for this oversight. We did not realize that during the pdf conversion process the bar corresponding to the DN-KASH group had changed to white. However, considering our reply to a concern raised by reviewer 1 (please see comments above), we have now decided to remove the section related to the chromosome segregation phenotypes and focus on the centrosome positioning mechanism instead.

*“Fig S6, albeit mentioned in the text, is not included in the supplementary info”.*

We apologize for this error. In fact, where it reads Fig. S6, should be Fig. S5. We have now corrected this.

*“a. GlutaMAX instead of GlutaMAXE (page 29)*
*b. What means "as described previously"? No reference is given. Do you refer to the upper part of the method section? (page 30)*
*c. 20 nM HEPES should most probably read 20 mM (page 32)*
*d. "1:50 protease inhibitor; 1:100 Phenylmethylsulfonul fluoride" - which protease inhibitor (mixture)? Rather phenylmethylsulfonyl fluoride.*
*e. exact composition of the cytoskeleton buffer used to prepare 4% paraformaldehyde could be given”.*

All these suggestions/corrections have been introduced in the text.

INSTITUTO
DE INVESTIGAÇÃO
E INOVAÇÃO
EM SAÚDE
UNIVERSIDADE
DO PORTO

Rua Alfredo Allen, 208
4200-135 Porto
Portugal
+351 220 408 800

www.i3s.up.pt
